# Graph-based pangenome of *Venturia inaequalis,* the apple scab fungus, reveals structural variants associated with population differentiation

**DOI:** 10.64898/2026.09.23.753852

**Authors:** Hana Feulner, Anze Svara, Xuebo Zhao, Zhangjun Fei, Awais Khan

## Abstract

Plant pathogen genomes adapt across coding and non-coding regions in response to their hosts, environments, and disease management practices. *Venturia inaequalis*, the causal agent of apple scab, has evolved alongside its *Malus* host throughout apple domestication. We generated 17 high-quality genome assemblies, including the first chromosome-scale assembly of *V. inaequalis*, to define the species’ complete gene content and investigate the role of structural variation in genome evolution. Together with a previously published scaffold-level reference, these assemblies were used to construct a gene-based pangenome containing 215,301 predicted protein-coding genes clustered into 13,240 orthogroups, 88.11% of which were core (73.86%) or soft-core (14.25%), 11.88% shell (present in 2-16 isolates), and 0.1% private, indicating a highly conserved gene repertoire. Virulence-associated effectors were significantly enriched in the soft-core and shell components, and accessory apoplastic effectors displayed elevated nucleotide diversity and nonsynonymous-to-synonymous substitution ratios (Ka/Ks) relative to core effectors, consistent with diversification. We further identified a putative accessory chromosome of variable size (23-132.6 kb) in European and U.S. isolates encoding genes with virulence function. A graph-based pangenome identified 38,799 non-redundant structural variants (SVs), largely driven by *Gypsy* LTR retrotransposons. We used the graph-based pangenome to genotype 136 globally distributed *V. inaequalis* isolates and identified variants differentiating populations associated with domesticated apple (*Malus domestica*) from those associated with its Central Asian wild progenitor, *M. sieversii*. Together, these resources establish a foundation for future studies of diversity, host adaptation, and genome evolution in *V. inaequalis*.

## Introduction

Apple scab, caused by the ascomycete fungal pathogen *Venturia inaequalis*, is among the most economically important fungal diseases of apples worldwide (MacHardy, 1996). The disease is found wherever apples are grown and is especially a major issue in temperate regions. Symptoms can occur on foliage and fruit, primarily affecting *Malus* species, although the pathogen can also infect other members of the *Rosaceae* family, including firethorn, mountain ash, and hawthorn (Jha et al., 2009). On leaves, the disease produces chlorosis, olive-green to gray, velvety lesions, and necrosis that can lead to defoliation and reduced vigor of the tree (Švara et al., 2024). On fruit, the pathogen causes corky, dark lesions that reduce marketability, resulting in substantial economic losses for growers (MacHardy, 1996). Fallen infected leaves and fruit serve as the primary source of inoculum and are the principal site of its annual sexual reproduction. Integrated management strategies include sanitation practices such as removing leaf litter to reduce primary inoculum and limit sexual reproduction of the fungus, as well as the use of resistant cultivars; yet chemical control remains the dominant management strategy (Nicholson and Rahe, 2004).

*V. inaequalis* and *Malus* exhibit a classic gene-for-gene interaction, in which individual resistance genes in *Malus* recognize corresponding fungal avirulence (*Avr*) genes, which typically encode small secreted proteins (Flor, 1971; MacHardy, 1996). To date, 18-19 gene-for-gene relationships have been described, with resistance genes termed *Rvi* followed by a number indicating the race specify of the pathogen and the order in which they were described (Bus et al., 2011; Patocchi et al., 2020). Despite extensive management efforts, the breakdown of host resistance conferred by the most widely deployed qualitative resistance alleles, including the most common scab resistance source, *Rvi6* from *Malus floribunda* 821, have been reported in orchards across Europe and the United States (Bengtsson et al., 1999; Papp et al., 2020b, 2020a; Patocchi et al., 2020).

These observations highlight the adaptive potential of *V. inaequalis*, consistent with patterns observed in other filamentous fungal plant pathogens that co-evolve with their hosts and adapt to changing environments under strong selective pressure. For *V. inaequalis*, this co-evolution has also occurred alongside its *Malus* host throughout the course of apple domestication (Ebrahimi et al., 2016; Gladieux et al., 2008). Modern cultivated apple is the product of genetic contributions from several wild *Malus* species, including *M. sieversii* from Central Asia, the primary progenitor of domesticated apple, as well as *M. sylvestris* from Europe, *M. orientalis* from Caucus, and *M. baccata* from Siberia (Cornille et al., 2014; Tegtmeier et al., 2025). Previous studies have identified genetically distinct populations of *V. inaequalis* associated with wild and domesticated apple hosts and demonstrated how gene flow between divergent pathogen populations inhabiting non-agricultural and agricultural hosts can maintain virulence alleles within populations (Gladieux et al., 2010; Lê Van et al., 2012).

Patterns of population differentiation and gene flow are shaped by ongoing host-pathogen co-evolution, which leaves detectable signatures of selection across pathogen genomes (Dong et al., 2016; Zande et al., 2023). In plant fungal pathogens, these signatures are often particularly evident in genes encoding small, secreted proteins known as effectors (Menardo et al., 2017). Effectors enable the pathogen to colonize the host by suppressing host immune responses and manipulating host physiology (Wu et al., 2022). However, some effectors can also be recognized by host resistance proteins, triggering immune responses that limit infection (Cook et al., 2015; Flor, 1971). This imposes strong selective pressures on pathogens to diversify their effector repertoires to maintain virulence (Sacristán et al., 2021). These adaptive processes can reshape pathogen genomes, resulting in gene gain and loss, structural variation, changes in ploidy, loss and gain of chromosomes, and the accumulation of repetitive elements (Galazka and Freitag, 2014; Kelkar and Ochman, 2012; Raffaele and Kamoun, 2012).

Earlier genomic studies of *V. inaequalis* identified an extensive effector repertoire, with effector genes frequently located near repetitive elements yet lacking the distinct “two-speed” genome organization observed in other filamentous pathogens where a repeat-poor, gene-dense core contrasts with a repeat-rich, fast-evolving accessory genome (Deng et al., 2017; Le Cam et al., 2019; Rocafort et al., 2022). Transcriptome-enabled studies have further characterized this repertoire, characterizing the infection process in temporal waves and finding many effectors in *V. inaequalis* have limited sequence homology to those in other plant pathogenic fungi yet have conserved structural features (Le Naour--Vernet et al., 2025; Rocafort et al., 2022). These include effectors in *V. inaequalis* with similar conserved domains to MAX effectors from *Magnaporthe oryzae (e.g., MoToxB, Avr-Pia)*, *Avr1* from *Fusarium oxysporum*, and Ecp10-like effectors from *Cladosporium fulvum (Mesarich et al., 2018; Rocafort et al., 2022; Seong and Krasileva, 2021; Yu et al., 2024, 2023)*. Recently, the first avirulence effector in *V. inaequalis* was identified and functionally validated. *AvrRvi6* was shown to adopt a MAX effector fold, similar to those described in the rice blast fungus *M. oryzae (Sannier et al., 2025)*. Multiple mechanisms enabling evasion of host immunity have been identified at the *AvrRvi6* locus, including mutations of key cysteine residues, a partial gene deletion, and a transposable element insertion in the promoter region (Lahfa et al., 2024; Sannier et al., 2025). However, these and other genomic studies of *V. inaequalis* have largely relied on short-read sequencing data or single, scaffold-level genome assemblies that were not fully resolved to chromosome scale, thereby limiting comprehensive genomics analyses of effectors across the species (Lichtner et al., 2020; Mellon et al., 2023; Papp et al., 2020b; Passey et al., 2018).

Advances in sequencing technologies have enabled the generation of numerous high-quality reference genomes, facilitating comparative analyses of genomic diversity and genome organization in fungal pathogens (Goodwin et al., 2011; van Westerhoven et al., 2025; Xu et al., 2025; Zaccaron et al., 2023). Studies across both plant and human systems have demonstrated that a single reference genome fails to capture the full extent of genetic diversity within a species or genus, giving rise to the concept of the pangenome–the complete genomic repertoire of a species (Li et al., 2023; Liao et al., 2023; Švara et al., 2024b). The pangenome comprises the core genome shared by all members of a species and the accessory genome that varies among individuals. In fungal plant pathogens, the accessory genome can contribute substantially to phenotypic diversity and is often enriched for genes associated with virulence and adaptation (Badet et al., 2020; McCarthy and Fitzpatrick, 2019; Tralamazza et al., 2024). Graph-based pangenomes further extend these analyses by integrating genetic variation across both coding and non-coding regions, including transposable elements, which can play critical roles in genome evolution, adaptation, and virulence (Faino et al., 2016; Torres et al., 2021). Graph-based pangenomes also enable the accurate detection of large structural variants (SVs), which are increasingly recognized as major contributors to genotypic and phenotypic diversity (Groza et al., 2024; Hartmann, 2022; Zhou et al., 2022).

In this study, we generated 17 high-quality *V. inaequalis* genome assemblies, including the first chromosome-scale reference assembly for the species. Comparative genomic analyses identified core genes within *V. inaequalis* and revealed effector genes exhibiting signatures of selection. Leveraging these assemblies, we constructed a graph-based pangenome to investigate the contributions of large SVs to genome evolution and their potential roles in geographic differentiation and host adaptation.

## Materials and Methods

### Isolate selection and DNA extraction

Single spore *V. inaequalis* isolates (**Supplementary Table 1**) were maintained on potato dextrose agar (PDA) supplemented with chloramphenicol (100 ug/mL) and streptomycin sulfate (100ug/mL). Isolates were obtained from apple-growing regions in the United States and Europe including historically relevant isolates such as those used in the PRI apple breeding program and INRAE isolates which have been characterized on the differential host set (Caffier et al., 2015). The collection also included isolates known to overcome *Rvi6* resistance gene, the most widely deployed apple scab resistance gene (**Supplementary Table 1**) (Belfanti et al., 2004). Mycelial plugs were transferred to 250-mL Erlenmeyer flasks containing 100 mL of potato dextrose broth (PDB) and incubated at room temperature with shaking at 90 rpm for approximately three weeks. Mycelium was harvested by vacuum filtration and lyophilized. High-molecular-weight (HMW) genomic DNA was extracted from 50 mg of lyophilized mycelial tissue using a modified cetyltrimethylammonium bromide (CTAB) protocol based on Schwessinger and McDonald (McDonald, 2017). Modifications included scaling down the input material, pooling multiple extractions from the same isolate to obtain a total DNA yield of 5 µg per isolate, and following DNA precipitation, DNA was spooled using a sterile glass rod, and then transferred to a clean tube prior to ethanol wash steps. DNA quantity and purity were assessed using the Qubit Broad Range dsDNA Assay Kit (Thermo Fisher Scientific) and a NanoDrop spectrophotometer (Thermo Fisher Scientific), respectively. DNA integrity was evaluated using pulsed-field gel electrophoresis on an Agilent Femto Pulse System at the Cornell Biotechnology Resource Center.

### Genome sequencing

Genomic DNA from all isolates was shipped on dry ice to the DNA Sequencing & Genotyping Center at the University of Delaware (USA) for Pacific Biosciences (PacBio) Single Molecule Real-Time (SMRT) sequencing. Genomic DNA was sheared to an average fragment size of 10-15 kb using a Megaruptor® 3 system (Diagenode), and libraries were prepared using the SMRTbell® Express Template Prep Kit v3.0 (PacBio). Barcoded libraries were pooled in equimolar amounts and size-selected using the BluePippin system (Sage Science) with a lower size cutoff of 6 kb. The pooled library was sequenced on a single SMRT Cell using the PacBio Revio™ sequencing system with Revio polymerase and sequencing kits, using a 30-h movie time.

Mycelium from isolate VI-19-031, collected in Geneva, New York from NovaEasygro, was prepared as described above and sent to CD Genomics (www.cd-genomics.com) for Hi-C library preparation according to previous studies (Padmarasu et al., 2019). Briefly, samples were cross-linked under vacuum infiltration with 3% formaldehyde for 30 min at 4℃ and quenched with 0.375 M final concentration glycine for 5 min. The cross-linked samples were subsequently lysed. After reversing cross-links, the ligated DNA was extracted using the QIAamp DNA Mini Kit (Qiagen) according to manufacturers’ instructions. Purified DNA was sheared to fragments of 300-500 bp, and were further blunt-end repaired, A-tailed and adaptor ligated, followed by purification through biotin-streptavidin-mediated pull-down and PCR amplification. Finally, the Hi-C libraries were quantified and sequenced on the Illumina Nova-seq platform (San Diego, CA, USA).

### Genome assembly

HiFi reads were used to generate 21-mer frequency distributions with Jellyfish v2.3.2 (Marçais and Kingsford, 2011), which were subsequently analyzed with GenomeScope2.0 (Ranallo-Benavidez et al., 2020) to estimate the genome size. HiFi reads were assembled into contigs using hifiasm v0.19.9 (Cheng et al., 2021) with the telomere motif-aware parameter ‘--telo-m GGGTTA’. Mitochondrial contigs were identified using MitoHiFi v3.0 (Uliano-Silva et al., 2023) and removed from the assemblies. Redundant or uncollapsed haplotigs were identified and removed using Purge Haplotigs (Roach et al., 2018). Quality of the assemblies before and after haplotig purging was assessed using QUAST v5.2.0 (Gurevich et al., 2013) to calculate contiguity statistics and completeness and Merqury v1.3 (Rhie et al., 2020) to assess base-level assembly accuracy and completeness. BUSCO v5.5.0 (Simão et al., 2015) was used with the *ascomycota_odb10* database to assess gene-space completeness. The assemblies were subsequently submitted to NCBI and screened for foreign contamination using the NCBI Foreign Contaminant Screening Pipeline v0.3.0 (O’Leary et al., 2024).

The assembly of isolate VI-19-031 was further scaffolded using Hi-C short-read data. Hi-C reads were aligned to the curated contig assembly using Juicer v2.0 (Durand et al., 2016b) and YaHS v1.2.2 (Zhou et al., 2023) was used to scaffold the contigs. The resulting assembly was visualized and manually curated using Juicebox Assembly Tools v2.17 (Durand et al., 2016a). The assembly was then finalized using the juicer post command and the liftover AGP file generated by YaHS, producing the final chromosome-scale assembly.

The 17 generated genome assemblies were anchored and oriented to the chromosomes of the newly generated chromosome-level assembly of VI-19-031 using RagTag v.2.0.1 (Alonge et al., 2022) with default parameters. Two assemblies containing chimeric contigs were corrected using ‘ragtag.py correct’ and validated using the original HiFi reads. For all assemblies, genome completeness was assessed again using BUSCO v5.5.0 with the *ascomycota_odb10* database. Consensus quality values (QV) and k-mer completeness were calculated using Merqury v1.3 (Rhie et al., 2020) to assess base-level assembly accuracy and completeness. Telomeric repeats (TTAGGG/CCCTAA) at both ends of chromosomes were identified using a modified version of the FindTelomeres.py script (Sperschneider, 2025).

### Genome annotation

Repeat libraries of interspersed repetitive elements were generated *de novo* for each genome using RepeatModeler v2.0.2 with the parameter “-*LTRStruct*” to enable the LTR structural discovery pipeline (Flynn et al., 2020). The resulting repeat libraries were then used with RepeatMasker v4.1.9 (http://www.repeatmasker.org/) to soft-mask the genomes in sensitive mode. RepeatMasker alignment output was processed using the parseRM.pl script, as described in Zaccaron et al., (2023), with the parameters “*--land* 50,1, *--parse*, and *--nrem*” to quantify the abundance of repetitive elements across different classes and families (Kapusta, 2025; Zaccaron et al., 2023). The same script was used to calculate the average percent divergence for each repeat family, and this information was then applied to estimate repeat divergence across the genome using a 20-kb sliding window. Composite RIP, repeat induced point mutation, index (CRI) for each annotated repeat by extracting the sequence from the genome and computing the transition from cytosine to thymine using the following formula, CRI = (TpA/ApT) − (CpA + TpG)/(ApC + GpT) (Selker et al., 2003).

Publicly available RNA-seq reads were downloaded from the NCBI Gene Expression Omnibus (accession GSE198244) (Rocafort et al., 2022) and used as transcript evidence for genome annotation. Raw paired-end RNA-seq reads were quality-trimmed using Trimmomatic v0.39 (Bolger et al., 2014). Trimmed reads were then mapped to each genome using HISAT2 v2.2.1 (Kim et al., 2019), and alignments with mapping quality scores (MAPQ) < 2 were filtered using SAMtools v2.0 (Li et al., 2009).

Protein-coding genes were predicted from the repeat-masked assemblies using the BRAKER3 pipeline (Gabriel et al., 2024). Transcript evidence was provided as described above, and protein homology evidence was obtained from the fungal protein dataset derived from OrthoDB (https://github.com/gatech-genemark/ProtHint#protein-database-preparation). BRAKER3 was run in ETP mode, in which GeneMark-ETP (Brůna et al., 2024) integrated both RNA-seq and protein homology evidence to generate initial gene predictions, followed by refinement with AUGUSTUS (Buchfink et al., 2015; Stanke et al., 2006). To improve annotation of effectors, two curated effector protein datasets (Rocafort et al., 2022; Sannier et al., 2025) were aligned to each genome using miniprot v0.13 with the options and filtering described in (Barragan et al., 2024, Li, 2023, https://github.com/YuSugihara/Barragan_and_Latorre_et_al_2024/tree/main).

Predicted proteomes and coding sequences were extracted using gffread v0.12.7 (Pertea and Pertea, 2020). For downstream comparative analyses, only the longest isoform per gene was retained using AGAT v1.2.0 (Dainat, 2026). Annotation completeness was then assessed using BUSCO with the ascomycota_odb10 lineage dataset prior to downstream analyses (Simão et al., 2015).

### Functional annotation and secreted protein annotation

Predicted proteins from VI-19-031 were functionally annotated by querying against the UniRef90 protein database using DIAMOND v2.0.9 (Buchfink et al., 2015) and the InterPro database using InterProScan v5.71 (Paysan-Lafosse et al., 2023). Gene Ontology (GO) terms were assigned using Blast2GO v1.5.1 (Conesa et al., 2005) by integrating homology-based evidence from DIAMOND with domain-based annotations from InterProScan. Secreted proteins were predicted for all isolates using SignalP v5.0 (Almagro Armenteros et al., 2019), and transmembrane domains were predicted using TMHMM v2.0 (Krogh et al., 2001). The secretome of each genome was then defined as proteins containing a predicted signal peptide and lacking transmembrane domains (Badet et al., 2020). Predicted secretomes were then analyzed using EffectP v3.0 (Sperschneider and Dodds, 2022) to identify proteins with putative effector function and predict their localization.

Effectors were assigned to expanded effector families from Rocafort *et al*. (2023), considering only families containing more than three members. For each family in Rocafort *et al*. (2023), protein sequences were aligned using MAFFT v7.520 (Katoh and Standley, 2013) with the “–auto” parameter. Alignments were trimmed using trimAl v1.4 (Capella-Gutiérrez et al., 2009) to remove poorly aligned and highly gapped regions, using a gap threshold of 0.3 and a minimum conservation threshold of 50%. Profile hidden Markov models (HMMs) were built from the trimmed alignments using hmmbuild in HMMER v3.4 (Finn et al., 2011) and searched against the *V. inaequalis* secretome using hmmsearch, with domain-level matches recorded using the “—domtblout” parameter. Each effector family was queried against the *V. inaequalis* secretomes using BLASTP v.2.16.0 (Altschul et al., 1997) with an E-value cutoff of 1e-5 and a minimum query coverage of 50%. HMMER hits were filtered based on domain-level significance (i-evalue < 1e-5). Results from BLASTP and HMMER searches were combined to assign proteins to effector families and the resulting gene assignments were mapped to their corresponding orthogroups.

### Gene-based pangenome and evolutionary analyses

Gene families across the 18 *V. inaequalis* assemblies, including 17 generated in this study and one previously published assembly (Le Cam et al., 2019), were identified using OrthoFinder v2.5.4 (Emms and Kelly, 2019), and implemented through GENESPACE v1.2.3 (Lovell et al., 2022). The gene-based pangenome was defined as the complete set of orthogroups identified across all genomes. Orthogroups were classified as core when present in all 18 isolates, soft-core when present in 17 isolates, shell when present in 2-16 isolates, and private when present in a single isolate. To model changes in pangenome and core-genome size with increasing numbers of genomes, pan- and core-genome sizes were calculated for all combinations of the 18 genomes at each genome count (N). Mean pan- and core-genome sizes at each value of N were used to generate accumulation curves. The pangenome accumulation curve was fitted using Heaps’ law, *P(N) = kN^α^*, whereas the core-genome curve was fitted using an exponential decay model (Tettelin et al., 2005).

The ratio of nonsynonymous substitutions per nonsynonymous site (*K_a_*) to synonymous substitutions per synonymous site (*K_s_*), *K_a_* /*K_s_*, was calculated for genes within each orthogroup. Gene sequences within each orthogroups were aligned using MAFT v7.520 (Katoh and Standley, 2013), and *K_a_* /*K_s_* values were calculated using KaKs_Calculator v2.0 (Wang et al., 2010). The median *K_a_*/*K_s_* value for each orthogroup was used for downstream analyses. Nucleotide diversity was calculated for each orthogroup using the ComputeStats function in EggLib v3.6.0 (Siol et al., 2022). Gene ontology (GO) enrichment analysis was performed using the topGO package (Alexa and Rahnenführer, 2024).

### Phylogeny and synteny analyses

Syntenic genomic regions among *V. inaequalis* assemblies were identified using McScanX v2.0 (Wang et al., 2012) and visualized using plot_riparian through GENESPACE v1.2.3 (Lovell et al., 2022). Gene-level synteny of chromosome 20 among *V. inaequalis* isolates was visualized using JCVI v.1.5.2 (Tang et al., 2024). BUSCO_phylogenomics v2023-12-17 was used to construct the alignment using single copy orthologs from the 18 *V. inaequalis* isolates in this study and *V. nashicola* (MAFF615029) (McGowen, 2023; Prokchorchik et al., 2019). A maximum-likelihood phylogeny was inferred using IQ-TREE v2.2. (Nguyen et al., 2015), with the best-fitting substitution model selected using the ModelFinder module and branch support assessed with 1,000 bootstrap replicates. The resulting phylogenetic tree was visualized using Interactive Tree Of Life (iTOL) (Letunic and Bork, 2021).

### Graph-based pangenome construction, SNP calling, and SV genotyping

The *V. inaequalis* graph-based pangenome was constructed of the 18 high-quality genome assemblies using the Minigraph-Cactus Package v2.6. (Wang et al., 2012), with isolate VI-19-031 used as the reference genome to initialize graph construction. The resulting pangenome graph was converted to a multi-sample VCF file, and overlapping and nested variant sites were normalized using vcfbub with the parameters ‘-l 0 -r 100000’ (https://github.com/pangenome/vcfbub).

SNPs and SVs were genotyped across 136 global *V. inaequalis* isolates from 16 countries across North America, Europe, Asia, and South America, using short-read resequencing data retrieved from the NCBI Sequence Read Archive (accession nos. PRJNA407103, PRJNA354841, PRJNA817384, and PRJNA962118) (**Supplementary Table 15**). Raw reads were processed using Trimmomatic v0.39 (Bolger et al., 2014) to remove Illumina adapter sequences and low-quality bases with the parameters “ILLUMINACLIP:2:30:10, LEADING:20, TRAILING:20, SLIDINGWINDOW:4:15, AVGQUAL:20, and MINLEN:25”. The resulting paired-end reads were then interleaved using SeqFu v1.10 (Telatin et al., 2021). SNPs and SVs were genotyped in each of the 136 isolates using PanGenie (Ebler et al., 2022). Variants with more than 20% missing genotypes were removed. Because *V. inaequalis* is haploid, genotypes in the graph-derived variant callset were represented as phased homozygous pseudodiploid genotypes to meet PanGenie input requirements. Multiallelic variant sites were decomposed into biallelic records using the PanGenie `convert-to-biallelic` script (https://github.com/eblerjana/pangenie/blob/master/pipelines/run-from-callset/scripts/convert-to-biallelic.py) prior to merging individual variant callsets across isolates with bcftools v1.20 (Li, 2011).

The resulting SNPs and SVs were processed and filtered using bcftools v1.20 (Li, 2011). Because *V. inaequalis* is haploid, heterozygous genotype calls were considered genotyping artifacts and masked as missing. Genotype calls with a genotype quality score below 15 (GQ < 15) were also masked as missing. Variant-level statistics, including minor allele frequency (MAF) and missing genotype rate, were calculated using the +fill-tags plugin in bcftools v1.20 (Li, 2011). Variants were retained if they had MAF ≥ 1% and missing genotype rate ≤ 20%. SNPs were extracted from the filtered variant callset and subjected to a more stringent missing genotype rate threshold of ≤ 10%. SVs were defined as insertions or deletions ≥ 50 bp in length.

SVs were functionally annotated using VEP v110.1 (McLaren et al., 2016) with the parameter ‘–pick’ to assign one annotation for each SV. To identify SVs associated with repetitive elements, alternative and reference allele sequences were extracted for insertions and deletions, respectively, using bcftools v1.20 (Li, 2011). These sequences were annotated using RepeatMasker v4.1.0 (Chen, 2004) against a classified repeat library generated from the reference isolate VI-19-031 using RepBase v26.04 (fngrep.ref) (Jurka et al., 2005). For SVs overlapping multiple repeat elements, the most abundant repeat class was assigned to the SV.

### Population genetic analysis

Principal component analysis (PCA) was performed separately for SNPs, insertions and deletions using PLINK v1.90b7 (Purcell et al., 2007). SNPs were converted to PLINK format and pruned for linkage disequilibrium using a sliding window approach with the parameters ‘`--indep-pairwise 50 10 0.2`’. PCA eigenvalues and eigenvectors for SNPs and SVs were visualized using ggplot2 v4.0.1 (Wickham, 2016). Pairwise genetic similarity was calculated separately from SNPs and SVs using the identity-by-state (IBS) metric implemented in PLINK v1.90b7 (Purcell et al., 2007). The resulting similarity matrixes were imported into R v2025.09.2+418 and converted to genetic distance matrixes. Neighbor-joining phylogenetic trees were constructed from the distance matrices using the ape package (Paradis et al., 2004) and visualized and annotated using the Interactive Tree of Life (iTOL) (Letunic and Bork, 2021). Population structure was inferred using ADMIXTURE v1.3.0 (Alexander and Lange, 2011) with SNPs pruned for LD as described above, and SVs that were pruned using PLINK v1.90b7 (Purcell et al., 2007) with the parameters ‘--indep-pairwise 500 50 0.5’. Two isolates, Vina 2493 from China and Vina 2508 from Japan, were excluded from downstream population analyses because their genetic clustering was inconsistent with their reported geographic origins. Nucleotide diversity (π) within each population and pairwise fixation index (*F*_ST_) between populations were calculated separately for SNPs and SVs using vcftools v0.1.16 (Danecek et al., 2011). Statistics were calculated both per site and in 20-kb sliding windows with a 10-kb step size.

SVs under selection were identified following (Zhao et al., 2026). Differences in SV allele frequencies between host-associated groups were assessed using Fisher’s exact test, with p-values corrected for multiple testing using the false discovery rate (FDR). SVs with FDR < 0.001 and a fold change > 2 were considered under selection.

## Results

### Chromosome scale assembly of a North American V. inaequalis isolate and 16 high-quality genome assemblies

We generated 8.86 Gb PacBio HiFi sequences for isolate VI-19-031, collected from NovaEasygro apples in Geneva, New York, with an average read length of 10,322 bp and corresponding to a genome coverage of ∼119x based on the estimated genome size of 74.35 Mb from k-mer analysis (**Supplementary Figure 1A**). The assembled contigs were scaffolded into chromosomes using 52.1 million paired-end Hi-C sequencing reads (**Supplementary Figure 1B**). The final VI-19-031 assembly contains 20 chromosomes, 18 contained telomeric repeats at both ends and two contained telomeric repeats at one end. The final assembly was 71.25 Mb, with an N50 value of 4.098 Mb, k-mer completeness rate of 99.30%, and a consensus quality value (QV) assessed by Merqury of 67.60 (**Supplementary Table 1**). BUSCO analysis using the ascomycota_odb10 indicated 98.0% completeness.

We also generated high-quality *de novo* assemblies of 16 additional *V. inaequalis* isolates using PacBio HiFi sequencing and included the previously published scaffold-level reference genome (Le Cam et al., 2019) to capture broader genetic diversity within the species (**Supplementary Table 1**).The average BUSCO of these 16 assemblies was 97.49%, and the mean consensus QV was 61.08. Telomeric repeats were detected at the ends of 313 of the scaffolds anchored to chromosomes, including 209 representing full chromosomes with telomeric repeats at both ends (**Supplementary Table 2**). The average N50 was 3.92 Mb and the assembled genome size ranged from 67.33 to 76.09 Mb (**Supplementary Table 1**). The number of predicted protein-coding genes ranged from 11,760 in VI-1771-2 to 12,363 in VI-EU301, and BUSCO completeness of the predicted proteins ranged from 94.20% in VI-1771-2 to 99.7% in VI-18-030 (**Supplementary Table 3**).

### Repeat landscape of V. inaequalis

The *de novo* repeat annotation of *V. inaequalis* isolates showed a similar repeat landscape. The number of repetitive bases ranges from 30.11 Mb in isolate VI-19-011 to 37.76 Mb (49.63%) in isolate VI-18-030, on average repeats covered 47.53% of the genome (**Supplementary Table 4**). Retroelements were the most abundant class of repetitive elements, accounting for 26.00-33.31% of individual genomes, whereas DNA transposons and unclassified repetitive elements accounted for 7.33% and 11.39 % of the genome, respectively. Long terminal repeat (LTR) retrotransposons were the predominant retroelement type, with Gypsy and Copia representing the major families (**Supplementary Table 4**).

Comparative analysis of repeats and their consensus sequences revealed that across all isolates, divergence profiles were broadly similar, with most repeats 75.47-82.18% of the total repeat content exhibited <10% divergence (**Figure 1A, Supplementary Figure 2A, Supplementary Table 5**). Divergence profiles did not differ significantly between European (n=7) and U.S. (n=10) isolates across any age class (**Supplementary Figure 3A**). Amongst all the genomes, Gypsy LTR retrotransposons contributed the largest proportion of the diverged TEs, followed by Copia LTR retrotransposons and unclassified elements (**Supplementary Table 6**).

**Figure 1.**
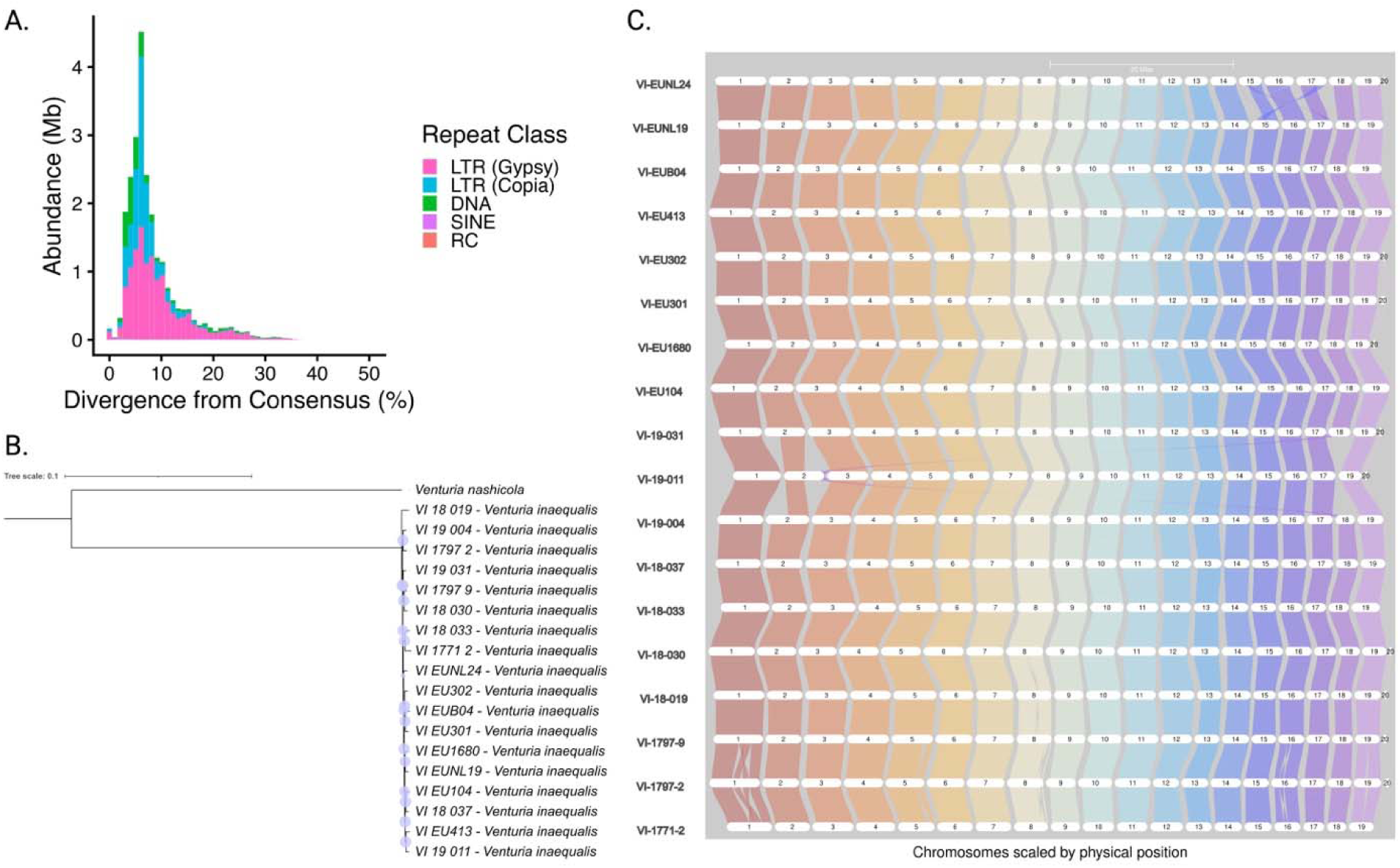
Repeat divergence, phylogenetic analysis and synteny analysis of *V. inaequalis* isolates. **A)** Bar plot showing the number of bases masked as transposable elements in *V. inaequalis* isolate VI-19-031, and TEs across classes and subclasses. The x-axis represents sequence divergence from the TE consensus, providing an estimate of age. **B)** Phylogenetic analysis of the of the 17 new *V. inaequalis* assemblies and the previous reference assembly ‘EU-B04’ and *Venturia nashicola* (Tree visualized using https://itol.embl.de/) using 1,658 single copy orthologs. Purple circles represent bootstrap values. **C)** Synteny map of the 17 new *V. inaequalis* genomes and previous reference (‘EU-B04’) based on gene order, using GENESPACE (https://github.com/jtlovell/GENESPACE). Shaded colors represent collinearity between regions.

Analysis of RIP (repeat-induced point mutation), a fungal genome defense mechanism that induces cytosine-to-thymine transition mutations in repetitive DNA, revealed that most TE families show evidence of RIP activity, with LTR retrotransposons being the most heavily affected class. Gypsy elements showed the highest and most consistent RIP signatures across all 17 isolates (mean CRI 1.44 ± 0.05, range 1.35-1.52), with 95.6% of elements having a CRI > 0. LTR Copia elements also showed strong RIP activity (mean CRI 1.05 ± 0.11, 87.3% CRI >0) (**Supplementary Figure 2B**). There was no significant different in RIP in TE families between the European and U.S. isolates (**Supplementary Figure 3B**).

### Phylogeny and synteny analysis

Phylogenetic analysis of *V. inaequalis* revealed low divergence among isolates, with U.S. isolates VI-19-011 and VI-18-037 clustering more closely with the European isolates (**Figure 1B**). Gene-based synteny analysis revealed broadly conserved macrosynteny across the genomes. However, small-scale rearrangements were detected, including those between isolates VI-EUNL24 and VI-EUNL19 on chromosomes 15, 16, and 17, primarily near the distal ends of these chromosomes (**Figure 1C**). Isolate VI-19-011, which had the lowest BUSCO completeness among the assemblies, lacked chromosome 18 but exhibited synteny between chromosome 2.

Additionally, a small putative accessory chromosome, chromosome 20, was identified in 13 isolates in both the European (n =4) and the U.S. populations (n=9) (**Supplementary Table 7, Supplementary Figure 4**). Ten of these isolates (n = 6 U.S and n =4 European) contained annotated genes on chromosome 20, comprising 17 orthogroups with no orthology to other chromosomes. Genes on this chromosome were associated with virulence including an apoplastic effector of unknown function, CAP10 domain-containing glycosyltransferase, a peptidyl-prolyl cis-trans isomerase involved in protein folding and secretion, and a SUN domain containing protein (**Figure 1B**). Additionally, a small putative accessory chromosome, chromosome 20, was identified in 13 isolates across both the European (n=4) and U.S. (n=9) populations (**Supplementary Table 7, Supplementary Figure 4**). Ten of these isolates (n=6 U.S., n=4 European) contained annotated genes on chromosome 20 comprising 17 orthogroups with no orthology to genes on core chromosomes; the remaining three U.S. isolates carried chromosome 20 but lacked annotated genes. Genes on this chromosome were associated with virulence, including an apoplastic effector of unknown function, a CAP10 domain-containing glycosyltransferase, a peptidyl-prolyl cis-trans isomerase involved in protein folding and secretion, and a SUN domain-containing protein.

### Gene-based pangenome

We used the 17 newly assembled *V. inaequalis* genomes together with the previously published reference assembly EU-B04 to construct and characterize the gene-based pangenome. Across the 18 genomes, 215,301 genes were assigned to 13,240 orthogroups (pangenes). Pangenes were classified into four categories: core, present in all isolates (73.86%; 159,029 genes); soft-core, present in 17 isolates (14.25%; 30,673 genes); shell, present in 2-16 isolates (11.88%; 24,572 genes); and private, present in a single isolate (0.01%; 27 genes) (**Figure 2A).** Core genes showed lower nucleotide diversity (π) and nonsynonymous-to-synonymous substitution ratios (K_a_/K_s_) than soft-core and shell genes (**Figure 2 C & D**). Pangenome size increased with the addition of each genome but began to reach a plateau after around 15 genomes. Fitting the accumulation curve using Heaps’ law yielded α = 0.0344, indicating that the pangenome remained open but expanded slowly with the addition of new genomes. The core genome decay curve approached a plateau, suggesting that the 18 genomes captured most of the core genes within the species (**Figure 2B).**

**Figure 2.**
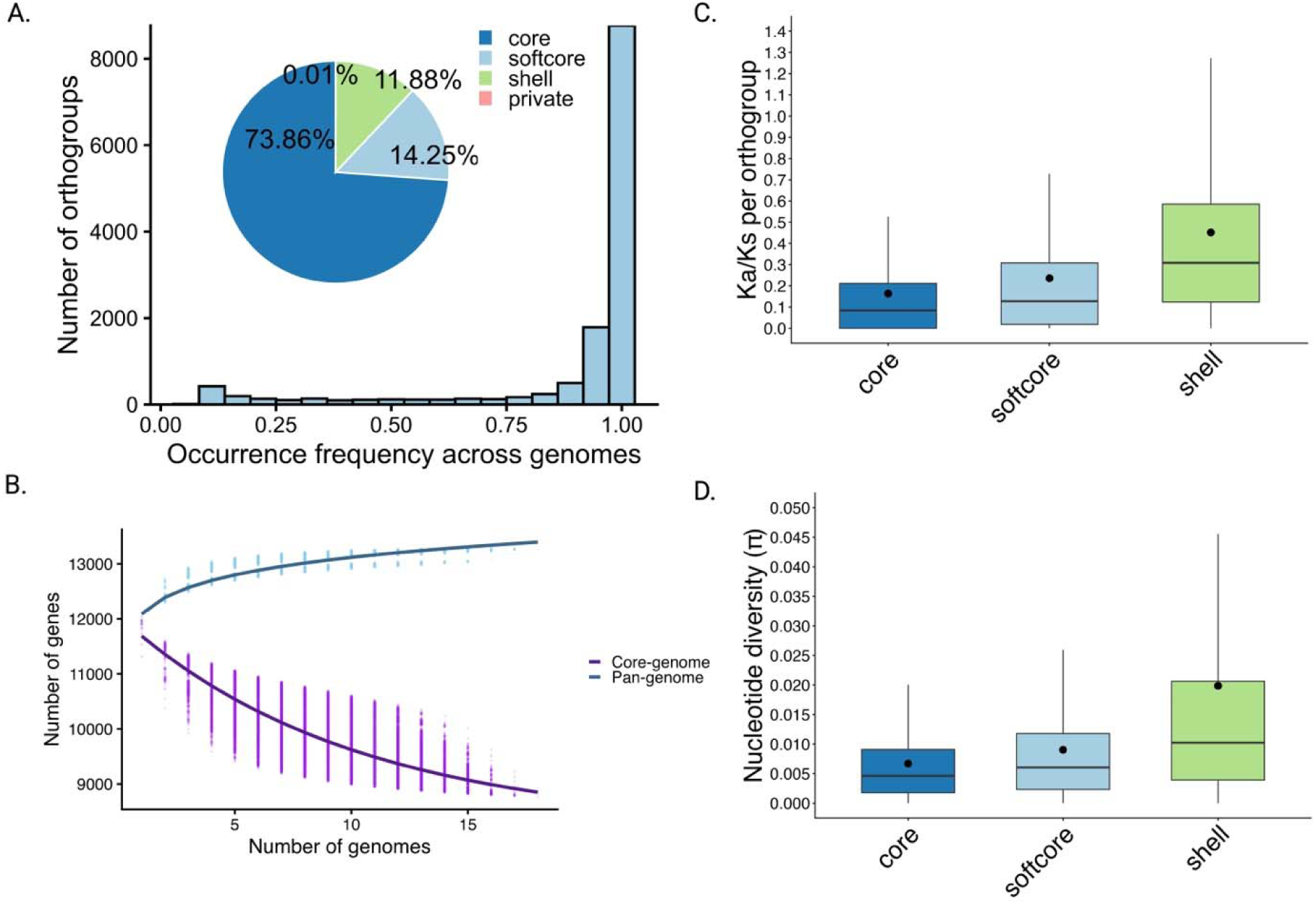
Gene based pangenome of *V. inaequalis*. **A)** Pie chart shows the composition of the gene based pangenome. The core represents genes found in all 18 *V. inaequalis* genomes, the softcore is present in 17 genomes, the shell represents genes in more than 2 but less than 16 genomes, and the private represents genes found in a single genome. The histogram shows the occurrence frequency of gene families (orthogroups) across the genomes. **B)** Pan and core genome modeling as additional genomes are added. Permutation-based pangenome (blue) and core genome (purple) sizes are shown with fitted curves. Light blue and purple points represent the observed pangenome and core genome sizes for each permutation, respectively. **C)** Median Ka/Ks per orthogroup for core (n=7,353), softcore (n=1,423), and shell (n=1,566). **D)** and Nucleotide diversity (π) for orthogroups. For each box plot, the lower and upper bounds indicate the first and third quartiles, respectively, the center line indicates the median, and the whiskers extend to 1.5× the interquartile range (IQR). Filled circles indicate group means.

GO enrichment analysis identified distinct functional profiles across pangenome compartments (**Supplementary Figure 4**). Core genes were enriched for fundamental cellular processes including metabolism, translational initiation, and meiosis (**Supplementary Figure 5A)**. Soft-core genes were enriched for processes such as phenylpropanoid metabolism and lipid transport (**Supplementary Figure 5B)**. Shell genes showed enrichment for terms associated with virulence and environmental adaptation, including chitin biosynthesis and transmembrane transport **Supplementary Figure 5C)**. The private genome consisted of 27 genes across 10 orthogroups distributed among seven isolates, most encoding hypothetical proteins of unknown function.

### Distribution and genetic diversity of effectors in the V. inaequalis pangenome

Across isolates, we identified between 1,588 and 2,109 secreted proteins, of which 700-1089 were predicted as candidate effectors by EffectP v3.0 (Sperschneider and Dodds, 2022) (**Supplementary Table 3**). At the pangenome level, we identified 710 orthogroups belonging to the expanded effector families described by Rocafort et al 2023. Of these orthogroups, 92% contained genes assigned to a single expanded effector family, whereas 8% contained genes assigned to multiple families (**Supplementary Table 8**). The largest family was effector family 1, a MAX like family, containing 1,333 genes, followed by family 3, a Ecp-10 like family containing 1,090 genes and the Nod19-family containing 735 genes across the isolates (**Supplementary Table 8**).

The distribution of effectors across pangenome categories revealed that 5.94% of the core genes were predicted to encode effectors, including 6,044 apoplastic and 3,407 cytoplasmic effectors. The proportion of predicted effectors increased to 13.04% in the soft-core (2,669 apoplastic and 1330 cytoplasmic), and 13.79% in the shell (2,460 apoplastic and 1,067 cytoplasmic) (**Figure 3A**). Effector genes were significantly enriched in soft-core and shell where they were 2.3- and 2.5-fold, respectively, more likely to occur compared to core genes (Fisher’s exact test, *p* < 0.001; **Figure 3B**). Most private genes encoded hypothetical proteins with unknown function, while two secreted proteins were identified in isolate VI-1771-2. These proteins encode identical apoplastic cysteine-rich secreted proteins, but lack recognizable conserved domains shared with other pathogenic fungi. The corresponding genes were located approximately 3.6 kb apart on chromosome 4, suggesting they may have arisen from a tandem duplication event.

**Figure 3.**
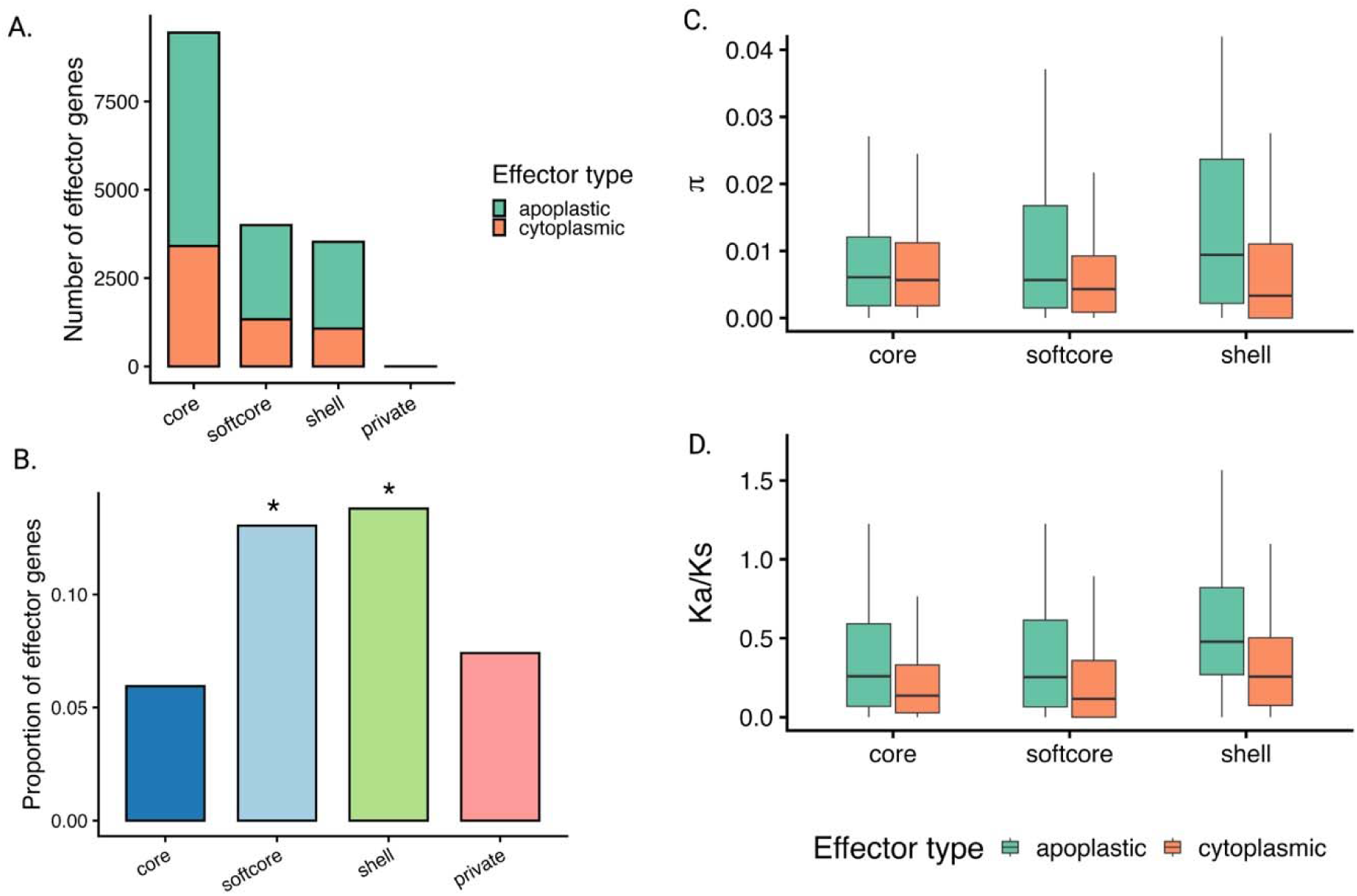
Effectors in the *V. inaequalis* pangenome. **A)** Distribution of genes annotated as effectors and their predicted localization across the core, shell, soft-core and private genomes. **B)** Proportion of effectors in each genome compartment, shell and softcore genomes were significantly enriched in effectors compared to the core genome (Fisher’s exact test, softcore *(p* < .05, Odds ratio 2.37) shell (*p* < .05, Odds ratio 2.53). **C)** Nucleotide diversity (π) and **D)** the ratio of nonsynonymous to synonymous substitutions (Ka/Ks) for single-copy apoplastic (apo) and cytoplasmic (cyto) effector orthogroups in core (apo: *n* = 300, cyto: *n* = 195), softcore (apo: *n* = 136, cyto: *n* = 80), and shell (apo: *n* = 194, cyto: *n* = 100) pangenome compartments. Boxes show the interquartile range with the median line; outliers are omitted for clarity. Statistical comparisons are reported in Supplementary Tables 9 and 11.

### Evolutionary analysis of cytoplasmic and apoplastic effectors in V. inaequalis

We next compared sequence diversity between predicted apoplastic and cytoplasmic effectors. Nucleotide diversity (π) did not differ significantly between apoplastic and cytoplasmic effectors in the core or soft-core. In the shell, however, apoplastic effectors exhibited significantly higher nucleotide diversity than cytoplasmic effectors (Wilcoxon test, adjusted *p* = 0.0004, **Supplementary Table 9, Figure 3C**). Among apoplastic effectors, nucleotide diversity was significantly higher in shell than those in core (adjusted *p* = 0.012), while no other pairwise comparisons among pangene categories were significant. Nucleotide diversity of cytoplasmic effectors did not differ significantly across pangene classes (**Supplementary Table 9, Figure 3C**). Core genes with the highest nucleotide diversity included a DUF3433-containing protein and six members of expanded effector families 3, 7, 29, and 68 among them MAX-like and Ecp10-like effectors and a chitin-binding type-4 domain-containing protein. Shell genes with the highest diversity included members of families 5, 11, 19, and 73, with families 3 and 19 encoding Ecp10-like effectors (**Supplementary Table 10**).

Apoplastic effectors exhibited significantly higher Ka/Ks values than cytoplasmic effectors across all pangenome compartments (core: adjusted *p* = 0.0021; softcore: adjusted *p* = 0.020; shell: adjusted *p* = 0.0021; Wilcoxon test, **Supplementary Table 11, Figure 3D**). Among apoplastic effectors, shell orthogroups showed significantly elevated Ka/Ks relative to both core (adjusted *p* < 0.001) and softcore (adjusted *p* < 0.001) genes, while core and softcore did not differ significantly from one another (adjusted *p* = 0.911). Cytoplasmic effectors followed a similar pattern but, with shell genes showing significantly higher Ka/Ks than core and softcore genes (adjusted *p* = 0.047 for both; **Supplementary Table 11**). Genes with Ka/Ks > 1, indicative of diversifying selection, many belonging to expanded effector families, were identified across pangenes (**Supplementary Table 12**): in the core genome, these included an Ecp10-like effector, a CFEM domain-containing protein, a MAX family effector, and members of families 2, 3, and 5; in the softcore, a MAX family effector, an ammonium transporter, and members of families 7, 11, 24, 28, and 35; and in the shell, a MAX-like effector and members of families 2, 3, 5, 12, and 35.

### Graph-based pangenome and identification of large structural variants

We constructed a graph-based pangenome using 18 high-quality *V. inaequalis* genome assemblies and identified 38,799 non-redundant large structural variants (variants larger than 50bp), comprising 13,418 deletions and 25,381 insertions. Collectively, these SVs represented 270,772,426 bp of cumulative sequence variation, with each additional genome contributing SVs gradually to the pangenome (**Figure 4A; Supplementary Table 13**). Most deletions (65.20%) were <1 kb, whereas 11.2% were >15 kb. Insertions showed a broader size distribution, with 49.4% <1 kb, 18.2% between 1 and 5 kb, and 14.3% >15 kb (**Supplementary Table 14; Supplementary Figure 6**).

**Figure 4.**
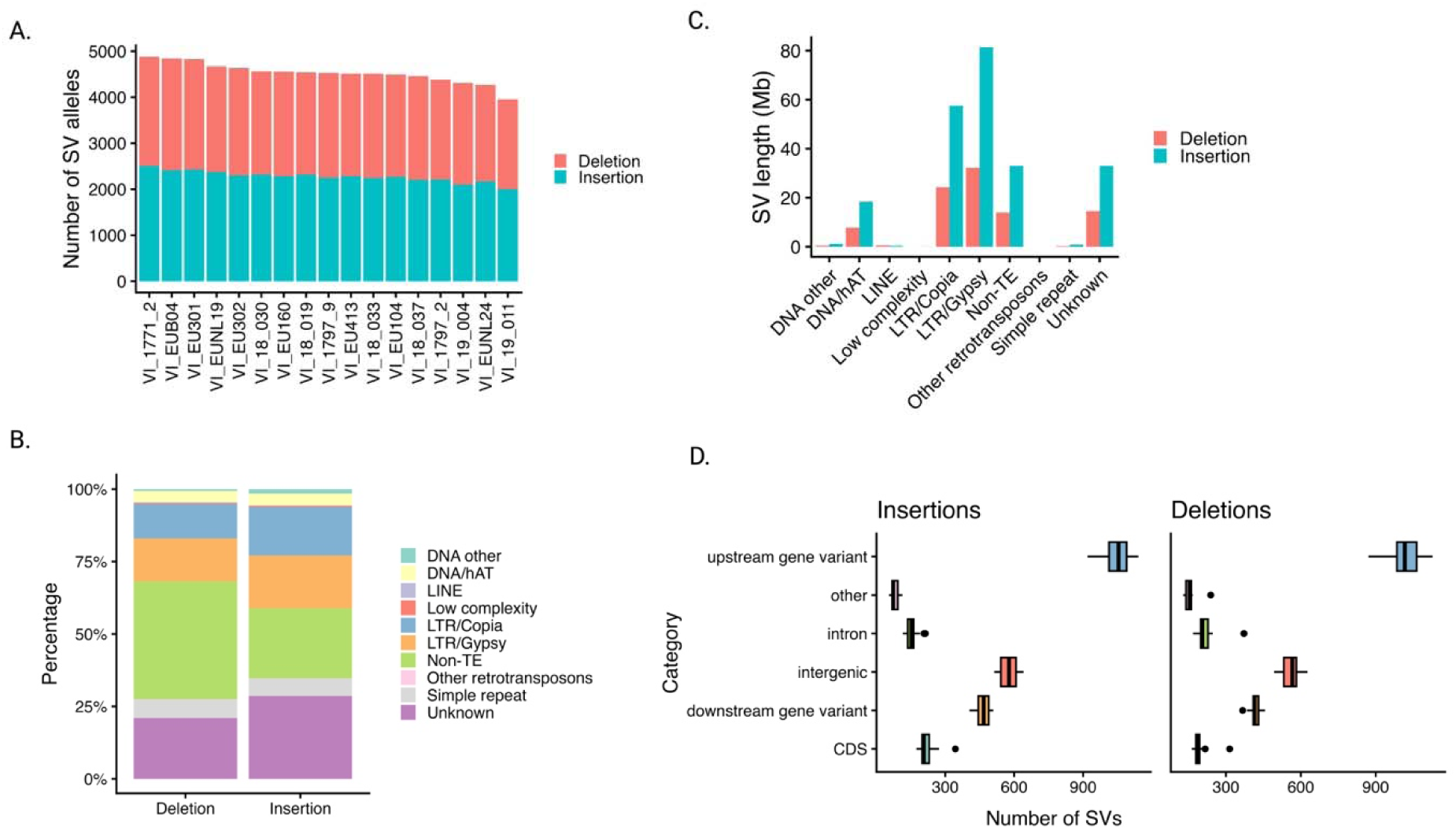
Structural variation in *V. inaequalis* isolates. **A)** Number of large structural variants (>50bp), colored by insertions and deletions in each isolate genome. **B)** Proportion of TE (transposable element) and non-transposable element derived structural variants, classified by the most common family found in the large structural variant. **C)** Total structural variant sequence (Mb) covered by transposable element families and non-TE families. **D)** Distribution of structural variant impacts across the isolates. Insertions left and deletions right. Downstream and upstream variants are within 1kb of genic regions.

Notably, 70.09% of SVs contained sequences derived from transposable elements (TEs). TE-associated sequences were detected in 59.28% of deletions and 75.81% of insertions. Among TE-associated SVs, unclassified repeat elements were the most abundant, accounting for 21.09% of deletions and 28.65% of insertions, followed by Gypsy LTR retrotransposons, which accounted for 14.77% and 18.17%, respectively (**Figure 4B**). TEs accounted for a large proportion of SV-associated sequence: 192 Mb of insertions and 79.7 Mb of deletions were TE-associated, compared with 46.9 Mb of non-TE sequence (33.0 Mb insertions, 13.9 Mb deletions; **Figure 4C**). Many SVs contained multiple repeat elements (**Supplementary Figure 7**). Most SVs occurred within 1 kb upstream of genes, followed by intergenic regions and regions 1 kb downstream of genes, whereas relatively few overlapped coding sequences. This pattern was consistent across insertions and deletions (**Figure 4D, Supplementary Table 15**).

### SV genotyping across isolates with distinct geography and hosts

We genotyped SVs represented in the pangenome graph in 136 publicly available resequenced *V. inaequalis* isolates using PanGenie v.4.21 (Ebler et al., 2022) to investigate their distribution across geography and hosts (**Supplementary Table 16**). A total of 32,020 SVs with <20% missing genotypes were retained for downstream population genomic analyses. The isolates are from a range of host species and geographic origins, with most from *Malus* hosts, including *M. domestica* (n = 51), *M. sieversii* (n = 21), *M. sieversii* and *M. domestica* (n = 20), *M. floribunda* (n = 8), *M. orientalis* (n = 6), *M. sylvestris* (n = 6), and *M. prunifolia* (n = 3). Non-*Malus* hosts included *Eriobotrya japonica* (n = 5), *Pyracantha* spp. (n = 5), and *Sorbus aucuparia* (n = 1), while host information was unknown for 10 isolates. The geographic origins included Central Asia, including isolates from the Central Asian Mountain (CAM), the Central Asian Plains (CAP), Western Asia (Armenia), Asia including China and Japan, the United States (US), and Europe (EU). Thirteen isolates were collected from hosts with *Rvi6*-mediated resistance. Phylogenetic analysis using SVs and SNPs clearly separated isolates based on their geographic origins and hosts (**Figure 5A, Supplementary Figure 8**). SV-based phylogeny resolved three major clades; isolates from Central Asia with a subclade of non-*Malus* host isolates, European isolates, and U.S. isolates. European and U.S. populations were clearly distinguished, within these populations, isolates collected from *Rvi6*-carrying hosts also clustered, a pattern also observed in the SNP-based phylogeny. The SNP-based phylogeny did not resolve European and U.S. populations as distinct clades, in contrast to the SV-based phylogeny (**Supplementary Figure 8**).

**Figure 5.**
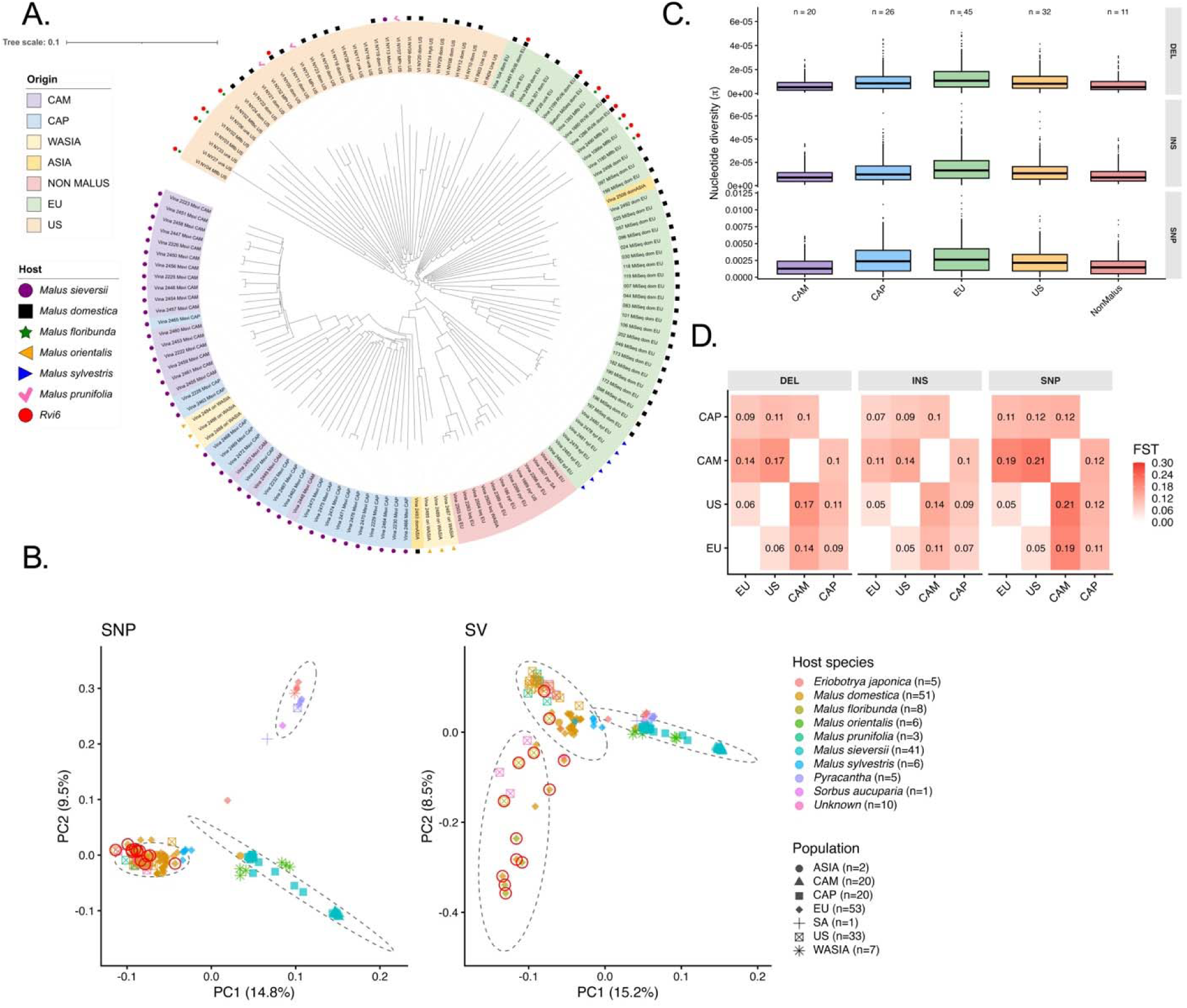
Population genetic analysis of *V. inaequalis* isolates using structural variants. **A)** Neighbor joining tree from large structural variants, rooted at midpoint. Colored labels indicate geographic origin or if isolates are from a non - *Malus* host (pink). Symbols represent *Malus* hosts if known and if an isolate was collected from a host with *Rvi6*. Tree visualized using iTOL (https://itol.embl.de/). **B)** Principal component analysis of 136 *V. inaequalis* isolates SNPs left and structural variants right. Dashed ellipses represent k-means clusters (k=3) identified from PC1 and PC2. Points are colored by the host they were collected from, shapes indicate geographic origin collected from, Red outline indicates the isolate was collected from an *Rvi6* host. **C)** Nucleotide diversity of deletions, insertions, and SNPs for the *V. inaequalis* populations, n indicates how many isolates were included in the population. **D)** F_st_ (allele fixation index) between the populations of *V. inaequalis* of deletion and insertion structural variants and SNPs.

Principal component analysis (PCA) and population structure analyses revealed marker type-dependent differences in isolate structure (**Figure 5B, Supplementary Figures 9-12**). K-means clustering (k=3) identified three groups in both SNP- and SV-based PCAs, though their composition differed. SNP-based PCA distinguished Central Asian isolates (CAM, CAP, WASIA, ASIA), European and U.S. isolates, and isolates from non-*Malus* hosts, with limited separation between European and U.S. populations. SV-based PCA instead grouped Central Asian and non-*Malus* isolates more closely while more clearly separating European from U.S. isolates. Interestingly, in the SV based PCA we observed distinct clustering of isolates from host with *Rvi6*, insertions also showed grouping of isolates from *Rvi6* hosts (**Supplementary Figure 9**). Population structure analyses showed a similar pattern where SV-based analysis showed admixture in the European population with components shared with Central Asian and U.S. populations, whereas SNP-based analyses showed less differentiation between European and U.S. populations (**Supplementary Figures 10 and 11**). Based on these analyses, *M. orientalis* isolates were grouped with the CAP population in downstream analysis.

The number and total length of SVs varied among isolates and populations, with European isolates carrying the most SVs and non-*Malus* hosts the fewest (**Supplementary Table 17, Supplementary Figure 13A &13B**). Insertions contributed a greater total sequence length across populations, reflecting a larger number of unique insertion events (**Supplementary Figure 14**). We next compared nucleotide diversity (π) within populations and pairwise fixation index (*F*_ST_) among populations using both SVs and SNPs; *F*_ST_ and nucleotide diversity estimated using SVs and SNPs were strongly correlated (*F*_ST_: r = 0.979-0.984; π: r = 0.954-0.964) (**Supplementary Table 18**). Nucleotide diversity was the highest and broadly similar in the European, U.S., and CAP populations, and lower in CAM and non-*Malus* host populations (**Figure 5C**). Pairwise *F*_ST_ revealed the greatest differentiation between U.S. and CAM populations, and between European and CAM populations (**Figure 5D**).

### SVs associated with geographic and host differentiation

To identify SVs associated with geographic and host differentiation, we compared SV allele frequencies between European and U.S. isolates and Central Asian populations (CAM and CAP), representing divergence associated with the geographic history of apple domestication. To minimize host-associated confounding effects, we restricted the European and U.S. populations only to isolates collected from *M. domestica*. A total of 1,026 significantly differentiated SVs were identified (false discovery rate < 0.001; fold-change > 2), including 302 enriched in European and U.S. populations and 724 enriched in isolates from Central Asia (**Figure 6A**). Of these SVs, 359 (35.0%) contained repeat sequences, while TE content did not differ significantly between populations (32.8% in European/U.S. and 35.9% in CAM/CAP). SVs enriched in European/U.S. isolates and CAM/CAP isolates impacted 244 and 544 genes, respectively, with 58 genes impacted by SVs enriched in both populations (**Supplementary Table 19**). Among genes impacted by differentiated SVs, 150 encoded predicted secreted proteins: 44 (18.3%) in European/U.S. and 106 (19.8%) in CAM/CAP.

**Figure 6.**
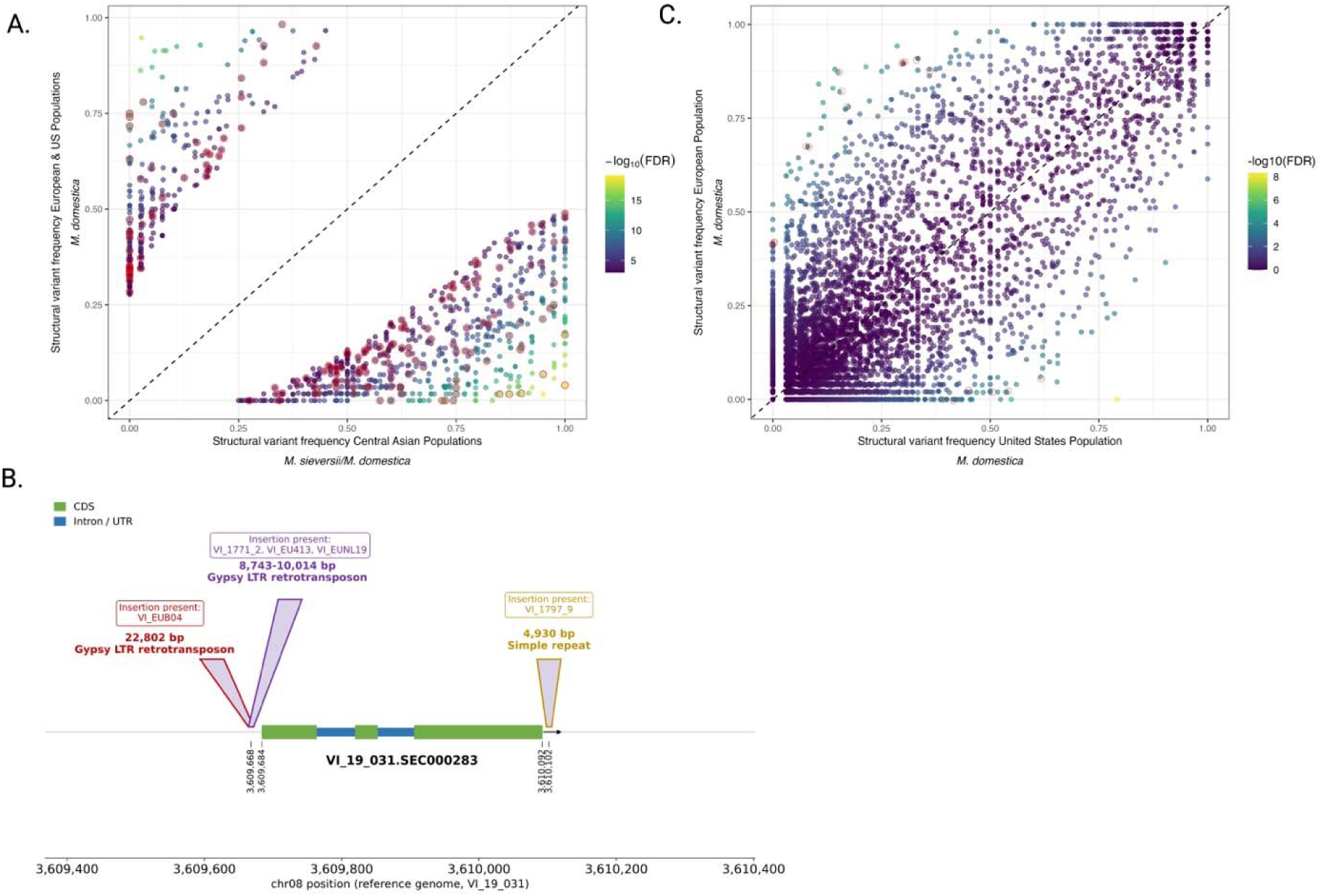
Structural variants under selection in *V. inaequalis* populations. **A)** European and United States populations compared to Central Asian populations. Top triangle shows SVs under selection in populations with from European and United States populations. Bottom triangle shows SVs under selection in central Asian populations (CAM and CAP). **B)** SVs at the VI_19_031.SEC000283 MAX-like effector locus on chromosome 8 on the ‘VI-19-031’ reference genome. The gene model is shown with CDS (green) and introns (blue); graph-level insertions within 2 kb are drawn at their corresponding reference breakpoints, labelled with insertion size, repeat class, and the assemblies carrying each. Two distinct LTR retrotransposon insertions (22,802 bp in ‘VI_EUB04’ and 8,743-10,014 bp in ‘VI-1771-2’,’VI-EU413’,’VI-EUNL19’) anchor 16 bp upstream of the transcription start, and a 4,930 bp simple-repeat insertion (‘VI_1797_9’) sits immediately downstream of the gene; SV is absent in CAM/CAP populations. **C)** United States compared to European isolates from *M. domestica*. Top triangle shows SVs under selection in isolates from Europe, bottom triangle shows SVs under selection in isolates from the United States.

We identified a 22.8-kb insertion affecting a MAX-like effector (VI-19-031.SEC000282) (**Supplementary Figure 15 A-C**) that was enriched in European/U.S. isolates (allele frequency = 43.9%) but absent from CAM/CAP isolates (FDR = 2.6 × 10⁻⁶, fold-change > 4 × 10⁵). The insertion was composed predominantly of Gypsy LTR retrotransposon sequence and was located 16 bp upstream of the annotated transcription start site, with predicted effects on the 5′ UTR and coding sequence. This is a site of recurrent, independent LTR retrotransposon insertion across the *V. inaequalis* pangenome. Of the assemblies in the graph, four (VI-1771-2, VI-EU413, VI-EUNL19 and VI-EUB04) carried the insertion composed almost entirely of Gypsy LTR retrotransposon sequence. In VI-1771-2, VI-EU413, and VI-EUNL19, the insertion ranged from 8.7 to10.0 kb and consisted predominantly of a single Gypsy element, with an additional ∼1.3-kb element nested within the 10.0 kb allele in VI-EUNL19. In VI-EUB04, the 22.8-kb insertion consisted of two distinct Gypsy retrotransposon families (**Figure 6B, Supplementary Figure 15 A**). The MAX-like effector was recovered in all assemblies except VI-1797-9, in which the syntenic region was assembled but masked by repeats. In each genome the respective insertions were confirmed to be upstream of the corresponding effector gene, suggesting a potential role for TE-associated structural variation in modulating effector expression.

We next directly compared European and U.S. populations and identified 89 differentiated SVs (FDR <0.001; >2-fold difference in allele frequency), of which 44 enriched in European isolates and 45 in U.S. isolates. Of the differentiated SVs, 48 (53.9%) contained repetitive sequence, with similar proportions among European- and U.S.-enriched SVs (54.5% and 53.3%, respectively; Fisher’s exact test, *p* = 1.0). These SVs affected 30 genes in EU-enriched set and 32 in US-enriched set, with two genes affected by distinct SVs in both sets (**Supplementary Table 20**). Among affected genes, 13 encoded predicted secreted proteins, including seven in the European-enriched set and 6 in the U.S.-enriched set, with no significant difference between populations (Fisher’s exact test, *p* = 0.76). Several highly differentiated SVs affected secreted genes with potential roles in host–pathogen interactions. A 939-bp insertion comprising 94% TE sequence occurred upstream of *g3819*, which encodes a predicted apoplastic effector with an AA1-like domain, and was enriched in U.S. isolates but absent from European isolates (allele frequency: U.S. = 0.39, European = 0.00; FDR 2.9×10^-⁴^). A 24.7-kb deletion upstream of a glycoside-hydrolase gene (*g10084*) was enriched in European isolates (European = 0.86, US = 0.33; FDR 7.7×10^-⁴^). We also detected a deletion of a complete gene encoding cytoplasmic effector (SEC002247). The gene was intact in most European isolates and deleted in approximately half of U.S. isolates (U.S. = 0.48, European = 0.02; FDR 9.3×10^-^⁵).

## Discussion

Studies across diverse biological systems, including humans, animals, plants and more recently fungi, have demonstrated that a single reference genome cannot capture the full extent of genetic diversity within a species (Azam et al., 2025; Gao et al., 2019; Liao et al., 2023; van Westerhoven et al., 2025b). Pangenome based approaches address this limitation by incorporating genomic diversity across multiple representative individuals. In particular, graph-based pangenomes, combined with long read sequencing technologies, can improve variant representation and detection especially for large SVs that are often missed using single references (Hickey et al., 2024). In fungal plant pathogens, recent studies have emphasized the contribution of SVs to genetic diversity and their potential roles in adaptive processes (Badet et al., 2020; Langner et al., 2021). In this study we generated 17 high-quality *V. inaequalis* genome assemblies, including the first chromosome-scale assembly for the species, enabling the construction of gene- and graph-based pangenomes that captured near-complete representation of the gene content and revealed extensive structural variation in the species, including 38,799 non-redundant SVs, many of which were associated with repetitive elements. Genotyping these variants across 136 globally distributed *V. inaequalis* isolates, including populations from Central Asia associated with *M. sieversii*, further associated SVs with geographic and host adaptation. These genomic resources provide a foundation for future comparative and functional genomic studies of *V. inaequalis* and its interaction with diverse host species.

We found that *V. inaequalis* has a largely conserved gene repertoire, with core genes accounting for 73.86% of the pangenome and a substantial soft-core component (14.25%) present in at least 17 of 18 isolates, together comprising 88.11% of the pangenome, with smaller shell (11.88%) and private (0.10%) components. Compared to other plant pathogenic fungi, *V. inaequalis* appears to have a relatively limited dispensable genome. For example, dispensable genomes of *Zymoseptoria tritici* and *Pyrenophora tritici-repentis* comprise approximately 45% and 57% of their respective pangenomes, respectively, whereas that of *Cladosporium fulvum* comprises approximately 2% (Gourlie et al., 2022; Wyka et al., 2022; Zaccaron and Stergiopoulos, 2024). The relatively conserved genome of *V. inaequalis* may be reflective of its unique infection strategy, it colonizes the host by using developing stroma and runner hyphae in the subcuticular space while not penetrating the plant epidermal cells, only later during infection are host cells (Bowen et al., 2011). It is possible the unique infection strategy may impose selection for a specific core gene repertoire, as the pathogen must sustain a compatible interaction with host tissue without triggering cell death. This study was also limited by the number of isolates and geographic regions sampled. Future studies incorporating a greater diversity of isolates from a broader range of locations would help validate and extend these findings.

Despite its relatively small accessory genome, we found that genes associated with virulence were significantly enriched in the soft-core and shell components of the *V. inaequalis* pangenome. Moreover, apoplastic effectors in the accessory components exhibited significantly higher nucleotide diversity and Ka/Ks relative to those in the core, indicating greater evolutionary variability among accessory effectors. Similar patterns have been reported in the grape ascomycete trunk pathogens *Eutypa lata* and *Phaeocremonium minimun*, in which dispensable biosynthetic gene clusters exhibited signatures of positive selection (Garcia et al., 2024). In *V. inaequalis*, amongst the dispensable genes with elevated Ka/Ks included additional members of previously described expanded effector families such as MAX-like effectors and numerous other families without structural homology (Rocafort et al., 2022).

Accessory chromosomes carrying genes related to virulence, host specificity and other adaptive processes have been described in many filamentous plant pathogenic fungi, including *F. oxysporum*, *M. oryazae* and *Zymoseptoria tritici* (Barragan et al., 2024; Habig et al., 2017; van Westerhoven et al., 2025). In *V. inaequalis*, we identified a small putative accessory chromosome ranging from 23 to 132.6 kb among a subset of both European and U.S. isolates. This chromosome contained 24 predicted secreted proteins classified as candidate effectors, 20 of which were not assigned to orthogroups containing effectors located on other chromosomes. BLASTp and HMMER searches revealed no detectable sequence homology between these proteins and characterized effectors in other plant pathogenic fungi, suggesting that they may represent highly diverged or *V. inaequalis*-specific effector candidates. We also identified a SUN domain-containing protein and a CAP10 domain-containing glycosyl transferase from the accessory genome. SUN domain-containing proteins have been implicated in reproductive development and host plant adhesion in *Botrytis cinerea* (Pérez-Hernández et al., 2017), whereas CAP10 has been identified as an important virulence factor implicated in polysaccharide capsule formation *Cryptococcus neoformans* and *Dactylellina haptotyla* (Tefsen et al., 2014; Wen et al., 2024). Further work will be required to elucidate the biological roles, distribution, and origin of this accessory chromosome, though its variable presence and enrichment for candidate effector genes suggest that it may contribute to phenotypic variation and geographic host adaptation in *V. inaequalis*.

The *V. inaequalis* genome is composed of, on average, 47.52% repetitive sequence, substantially higher than other members of the genus, which range from 2.5% in *V. nashicola* to 17.61% in *V. effusa* providing evidence for a genome which has undergone expansion due to repetitive elements (Prokchorchik et al., 2019; Winter et al., 2020). Importantly, these genomes were assembled using third-generation long-read sequencing, making it unlikely that their lower repeat content reflects assembly gaps rather than true biological differences. The TE families in *V. inaequalis* in this study overall have low repeat divergence indicating on-going expansion of transposable elements. Interestingly in this study and previous studies found evidence of repeat induced point mutation (RIP), a genome-defense mechanism expected to increase sequence divergence among repetitive elements, which occurs before meiosis, (Galagan and Selker, 2004; Le Cam et al., 2019). The low sequence divergence and detection of RIP may be an effect of ongoing TE expansion that is actively being countered by genome defense mechanisms, though this is unclear. A similar pattern has been reported in *Cladiosporum fulvum*, in which low repeat divergence was interpreted as evidence of recent TE proliferation, yet extensive RIP was identified. This pattern was proposed to result from infrequent sexual reproduction limiting the accumulation of RIPs (Zaccaron and Stergiopoulos, 2024). Further, we did not observe any significant differences in divergence of TE families or RIP between the European and United States populations, despite hypothesizing that we might see evidence of more recent expansion and less RIP in the United States population, given that those isolates were collected more recently and are derived from the European population.

Structural variation in *V. inaequalis* has remained largely unexplored. Here, we used a graph-based pangenome approach to identify and characterize large SVs, many of which were associated with TEs. Among TE-associated SVs, Gypsy LTR retrotransposons contribute the largest fraction and were also among the most abundant classes in the genome. Most insertions and deletions occurred within 1 kb upstream of genes, placing substantial structural variation in regions with the potential to influence gene regulation. Previous studies in *V. inaequalis* have similarly shown that candidate effector genes tend to occur in close proximity to repeat elements (Palacıoğlu et al., 2022; Tenzer and Gessler, 1999). TE mobilization has been shown to affect virulence in fungal pathogens, such as in *V. inaequalis* TE insertion causes disruption of the *AvrRvi6* effector, enabling isolates with this insertion to overcome *Rvi6*-mediated host resistance (Kang et al., 2001; Sannier et al., 2025; Wu et al., 2015). These findings highlight the potential for TE-associated structural variation to contribute to the evolution and regulation of genes in *V. inaequalis*. Future studies will be needed to determine whether the TE-associated SVs identified here, specifically population-differentiated variants affecting candidate effectors, modulate expression or function of specific effectors and effector families in *V. inaequalis*.

Genotyping of variants across 136 globally distributed isolates revealed that SVs were associated with both geographic origin and host. Population structure inferred from SNPs (*K* = 3) was consistent with previous findings, showing limited differentiation between European and U.S. populations (Mellon et al., 2023). In contrast, SVs provided greater resolution between these populations, although admixture was still observed. This suggests that structural variation captures population differentiation not captured by SNPs alone and highlights the power of a graph-based pangenome in characterizing population-level structural variation (Du et al., 2025). A limitation of this study, however, is the lack of representation of Central Asian isolates in the pangenome graph, restricting genotyping to variants only already present in the graph (Ebler et al., 2022). In plants, inclusion of diverse accessions representing cultivars, landraces and wild relatives in pangenome construction has facilitated the detections of SVs associated with domestication and adaptation (Sun et al., 2025; Zhao et al., 2026). Similarly, future studies in *V. inaequalis* might include isolates from more diverse geography.

Despite this limitation, the presence of shared SVs across Central Asian, European and U.S. populations enabled the identification of variants showing strong allele-frequency differentiation among these *populations.* Previous studies have identified *M. sieversii* as the ancestral host of *V. inaequalis* and have shown that apple domestication has shaped pathogen genetic diversity, resulting in high within-population diversity and relatively low geographic differentiation (Ebrahimi et al., 2016; Gladieux et al., 2010). Consistent with this, our results demonstrate that SVs shared across populations can still capture signatures of host-associated adaptation between European and U.S. populations associated with *M. domestica* and Central Asian populations associated predominantly with *M. sieversii*. Notably, many of the genes affected by these differentiated SVs encoded predicted secreted proteins with putative roles in host-pathogen interactions. These findings highlight the importance of structural variation in shaping host adaptation during the spread of *V. inaequalis* from Central Asia and wild *Malus* progenitors to cultivated apple.

## Conclusion

Here, we generated genomic resources that expand the foundation for comparative and population genomics of *V. inaequalis*, one of the most important fungal pathogens of apples. We demonstrate the value of pangenomic approaches in fungal genomics by moving beyond a single reference genome to characterize gene content and structural variation within the species. We found in *V. inaequalis* gene content is conserved with core and soft-core genes together making up 88.11% of the pangenome (73.86% and 14.25%, respectively). Virulence-associated genes were enriched in the soft-core and shell genome, and accessory apoplastic effectors displayed significantly higher nucleotide diversity and Ka/Ks ratios than core effectors, indicating these may be under diversification. We found extensive structural variation using a graph based pangenomic approach, identifying 38,799 non-redundant SVs, largely driven by *Gypsy* LTR retrotransposons. Across 136 globally distributed isolates, these SVs provided greater resolution of European and U.S. population structure than SNPs and identified variants differentiating populations associated with cultivated *M. domestica* from its Central Asian progenitor *M. sieversii*. We further identified a putative accessory chromosome (23-132.6 kb) in both European and U.S. isolates that encodes punitive virulence genes. Future work, expanding on these resources, including additional long read assemblies from diverse geographic and host populations, will enable pangenome-wide association studies linking phenotypes to genotypes in this pathogen and other economically important fungal pathogens,

## Data availability statement

The datasets supporting the conclusions of this article comprising HiFi and Hi-C raw reads has been deposited in the NCBI under the Bioproject accession number PRJNA1180949.

## Declarations

Not applicable

## Code Availability

Code for this analysis is available at https://github.com/hana-a-f/V.-inaequalis-pangenome

## Acknowledgments

We acknowledge the help of Della Cobb-Smith for fungal sample preparation for DNA extraction and sequencing. We also acknowledge the help of Dr. Qi Sun for IT support during data analysis, Dr. Jacob Landis for his support during Hi-C data analysis.

## Author Contributions

H.F. and A.K. designed the experiment. H.F performed the data analysis. X.Z. assisted in phylogenetic analysis and graph based pangenome analysis. A.K. supervised the research and secured funding. H.F. wrote the manuscript. A.K, A.S. and Z.F. revised the manuscript. All authors have read and approved the manuscript.

