## Supplementary Figure for "Graph-based pangenome of *Venturia inaequalis,* the apple scab fungus, reveals structural variants associated with population differentiation"

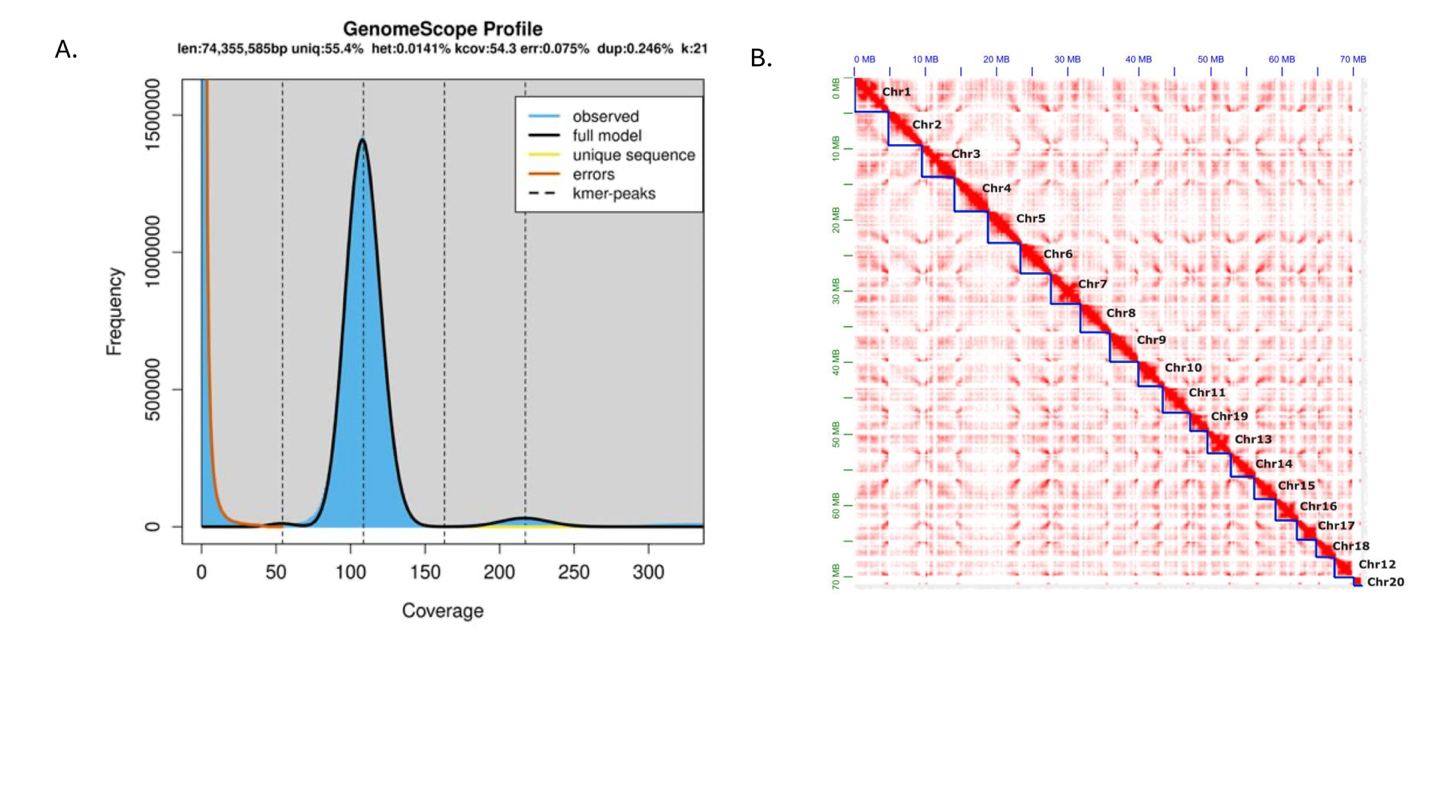

**Supplementary Figure 1**. K-mer frequency distrubtion and Hi -C interaction map for chromosome scale assembly of *Venturia inaequalis* (VI-19-031). **A)** k-mer frequency distribution of *V. inaequalis* isolate VI-19-031. The GenomeScope profile shows a single major k-mer peak at ~119× coverage, consistent with a haploid genome, estimated genome size 72 Mb with 0.03 % heterozygosity and 3.2 % duplication. **B)** Chromatin contact map of *Venturia inaequalis* isolate ‘VI-19-031’. Chromosome-scale Hi-C interaction map showing contact frequency across the 20 chromosomes of the haploid *V.* *inaequalis* assembly.

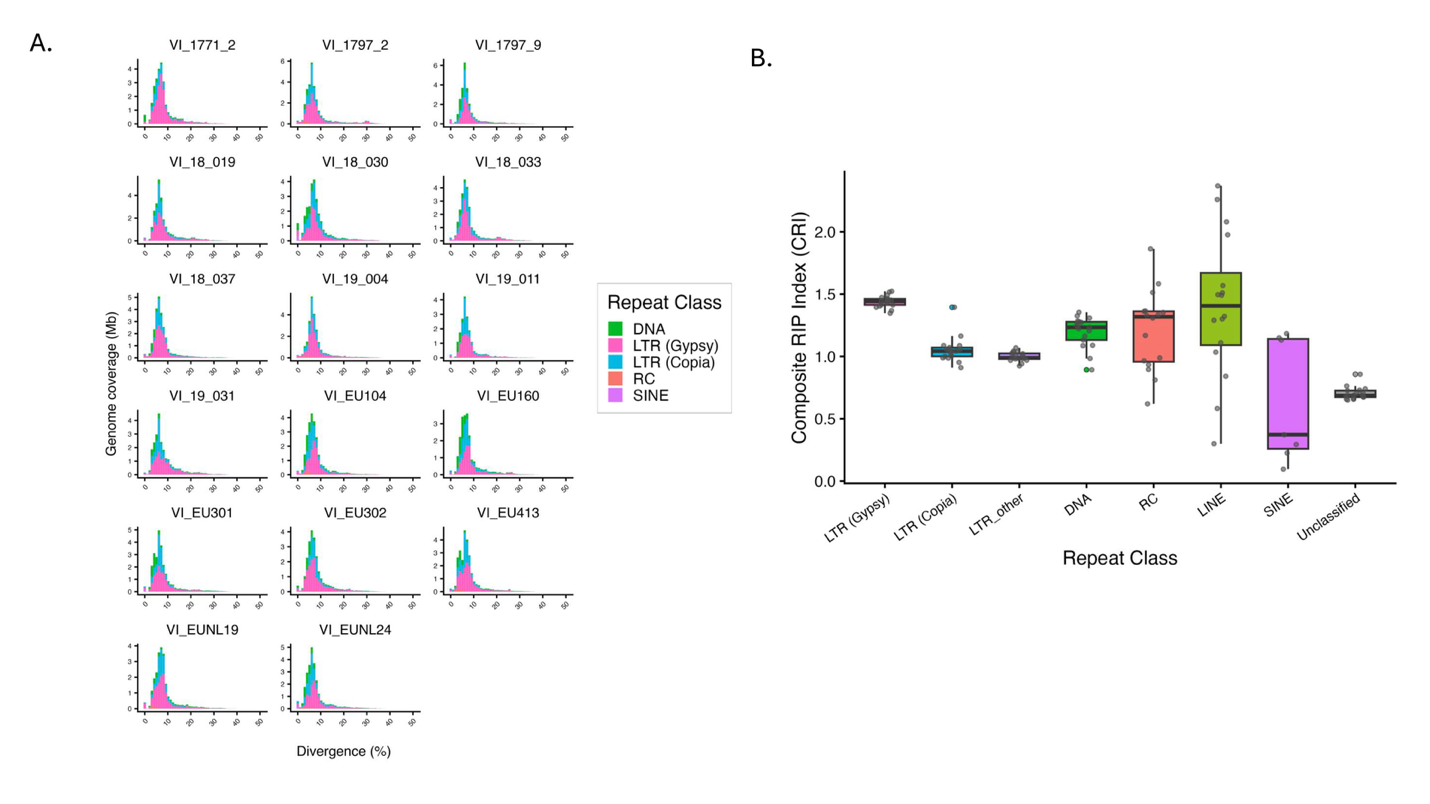

**Supplementary Figure 2.** Repeat divergence and repeat induced point mutations of transposable element families in *V. inaequalis****.* A)** Bar plot showing the number of bases masked as transposable elements (TEs) across classes and subclasses (LTR Gypsy/Copia); DNA transposons (DNA), long interspersed nuclear elements (LINE), long terminal repeats (LTR), rolling-circle elements (RC), and unclassified repeats across *V. inaequalis* isolates. The x-axis represents sequence divergence from the TE consensus, providing an estimate of age. **B)** RIP activity across transposable element families in *V. inaequalis*. Composite RIP Index (CRI) calculated per TE copy and summarized across 17 new isolates. Each point represents the mean CRI for one TE family in one isolate; boxes show the median and interquartile range.

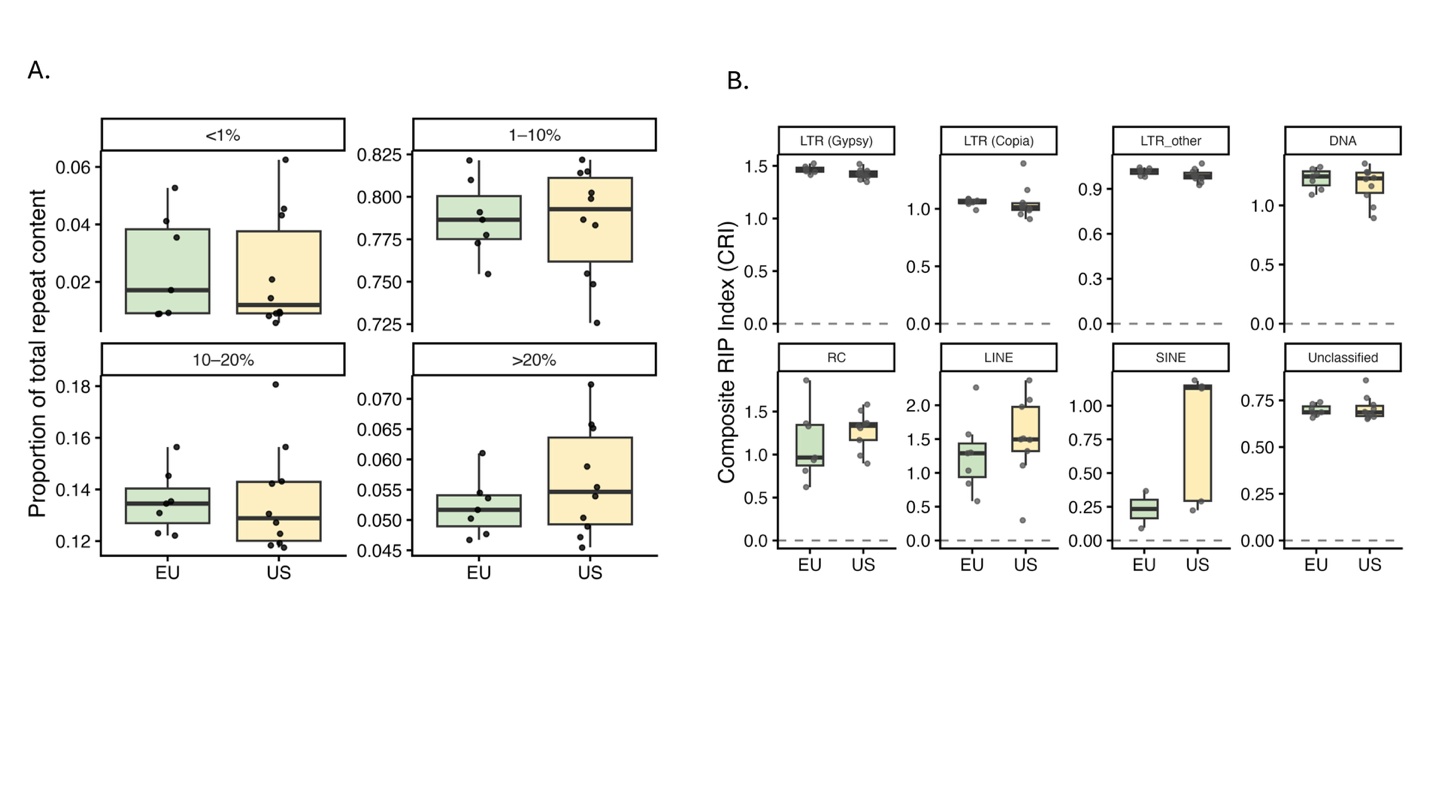

**Supplementary Figure 3**. Comparison of transposable element divergence and composite RIP index (CRI) between European (n=7) and United States (n=10) *V. inaequalis* isolates. Box plots show median, interquartile range and 1.5× IQR. **A)** Proportion of total repeat content within each divergence class (<1%, 1-10%, 10-20%, >20%). No significant differences were detected between populations for any divergence class (two-sided Wilcoxon rank-sum test with Benjamini-Hochberg correction; <1%: W=37, *p*=0.887; 1–10%: W=33, *p*=0.887; 10–20%: W=41, *p*=0.601; >20%: W=26, *p*=0.417; all *p* adj =0.887). **B)** Comparison of Composite RIP Index (CRI) per TE family. No significant differences were detected between populations for any TE family (two-sided Wilcoxon rank-sum test with Benjamini-Hochberg correction; LTR Gypsy: W=56, *p*=0.043, *p* adj =0.345; LTR other: W=51, *p*=0.133, *p* adj=0.430; LTR Copia: W=50, *p*=0.161, *p* adj =0.430; RC: W=21, *p*=0.299, *p* adj=0.479; LINE: W=21, *p*=0.299, *p* adj =0.479; SINE: W=2, *p*=0.381, *p* adj =0.508; DNA: W=40, *p*=0.669, *p* adj=0.765; Unclassified: W=37, *p*=0.887, *p* adj =0.887).

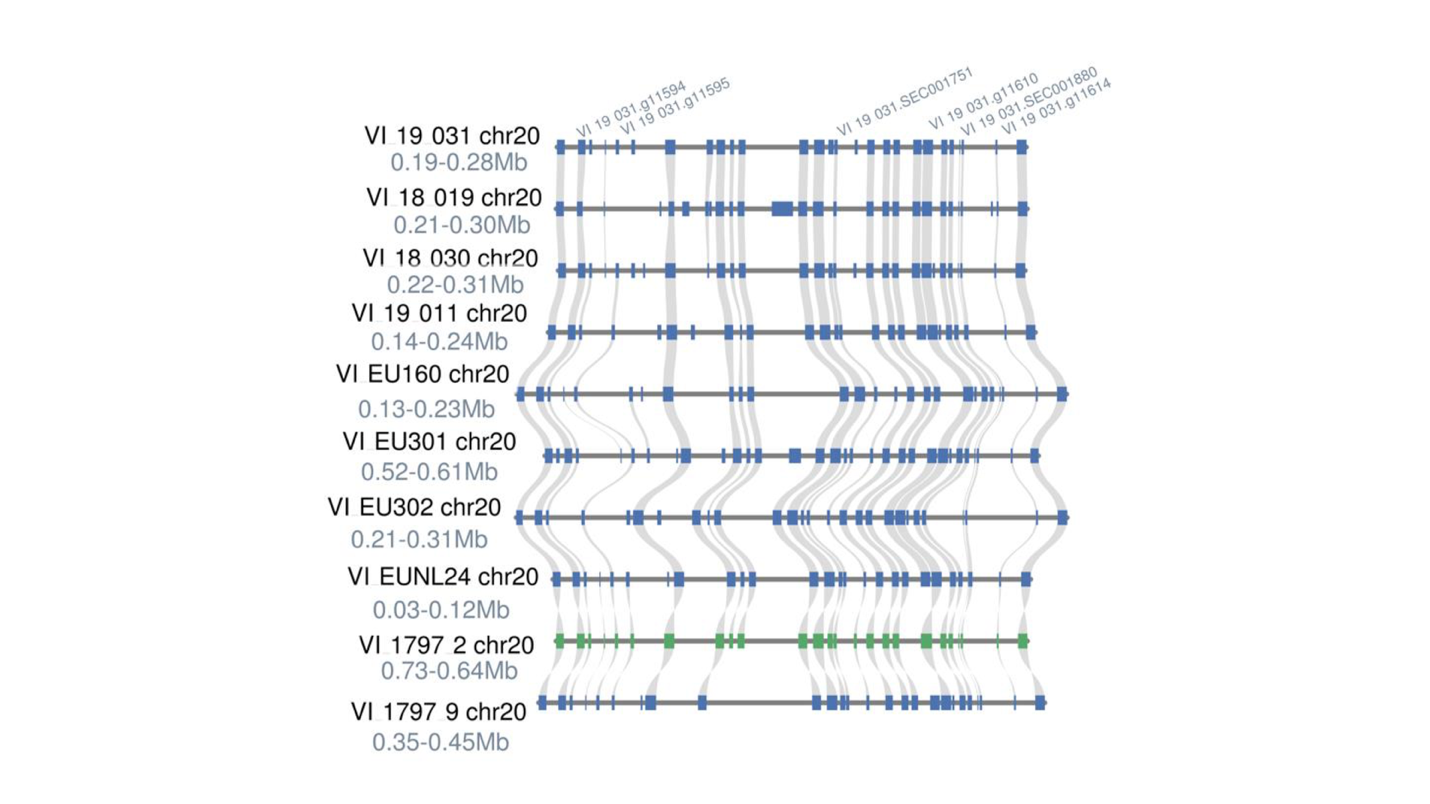

**Supplementary Figure 4.** Accessory chromosome of *V. inaequalis*. Synteny comparison of chromosome 20, genes with punitive fitness advantage are shown in text (VI-19-031.g11594: Peptidyl-prolyl cis-trans isomerase,VI-19-031.g11595: CENP-A homolog, VI-19-031. SEC001751: cytoplasmic effector, VI-19-031.g11610: SUN domain-containing protein, VI-19-031.SEC001880: apoplastic effector, VI-19-031.g11614: Glycosyl transferase CAP10 domain-containing protein), green color of VI-1792-2 indicates the sequence is inverted compared to the other isolates. All genes are listed in Supplementary Table 8.

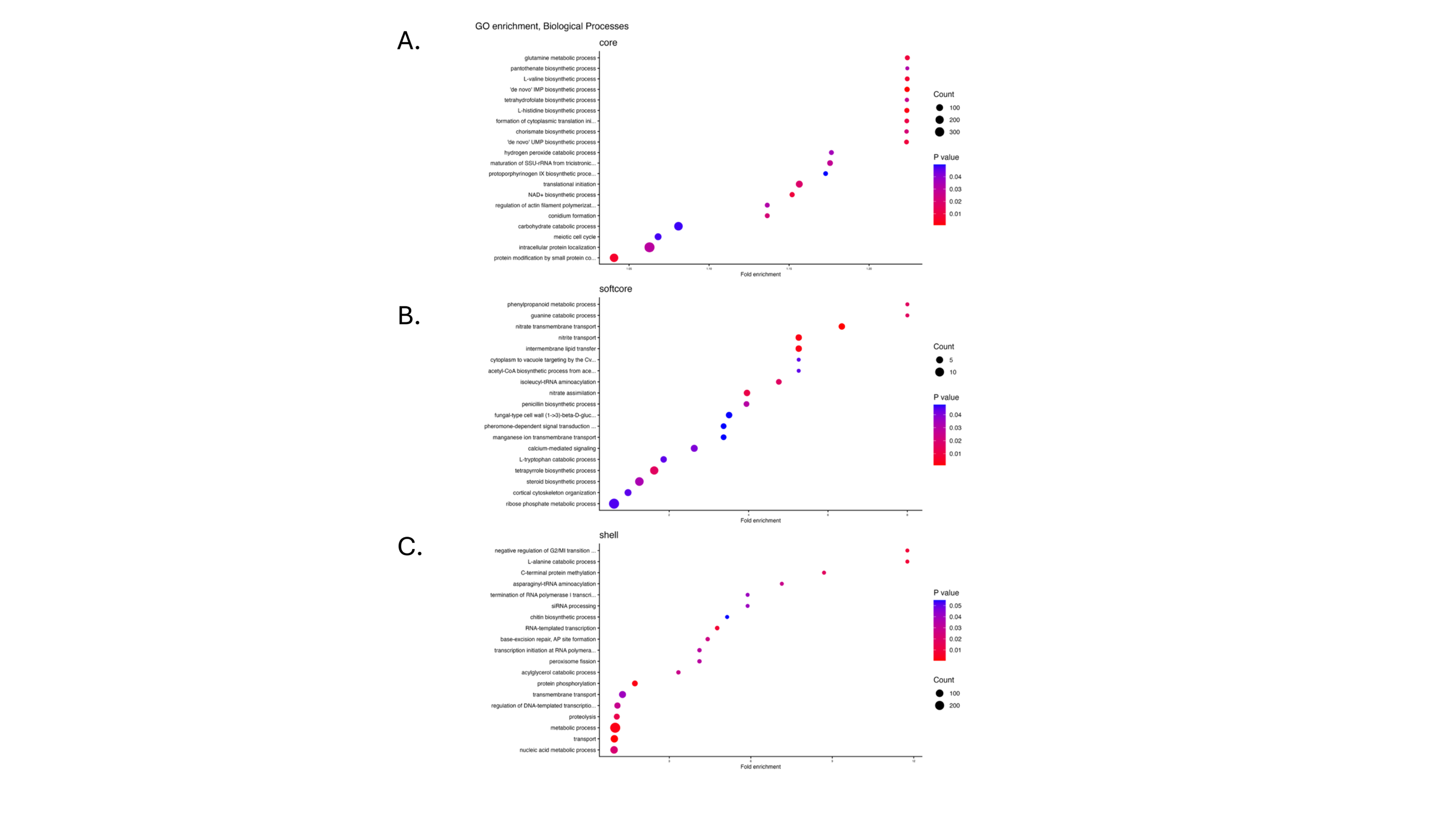

**Supplementary Figure 5.** GO enrichment of *V. inaequalis* pangenome. Functional enrichment of biological processes of the **A)** core, **B)** softcore and **C)** shell genes in the *V. inaequalis* gene based pangenome.

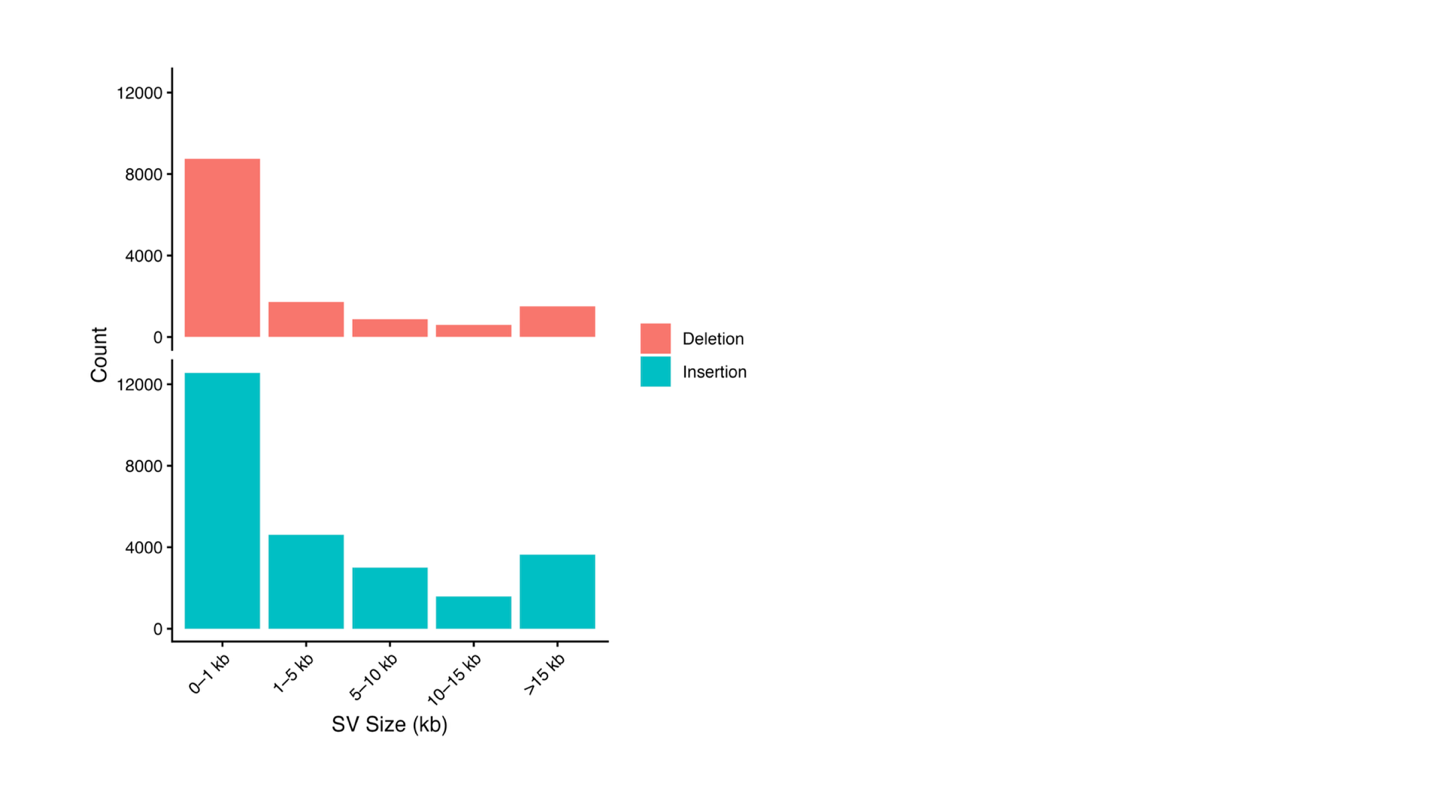

**Supplementary Figure 6.** Distribution of SV length (kb) in *V. inaequalis*. Top shows deletions, bottom shows insertions.

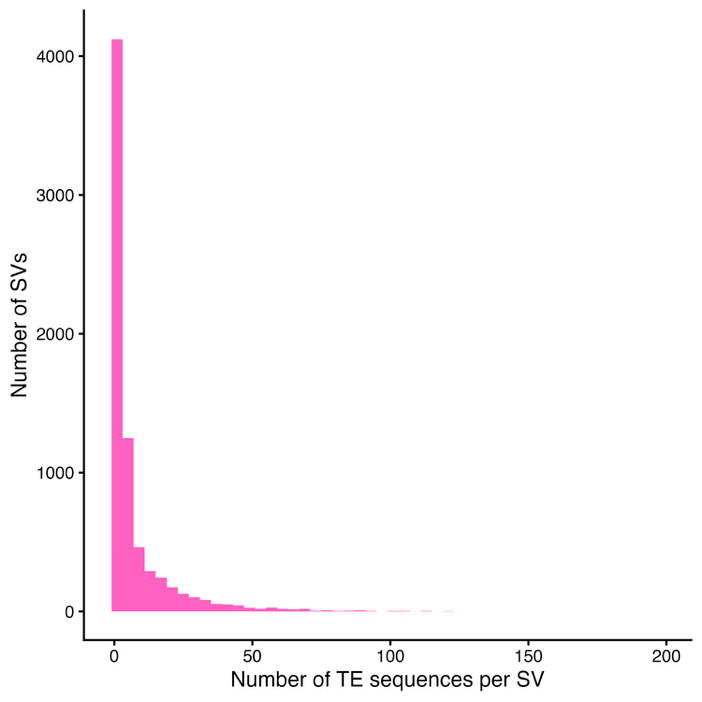

**Supplementary Figure 7**. Structural variants containing repeat elements. The x-axis shows the number of TE sequences identified in a single SV, y-axis shows the number of SVs.

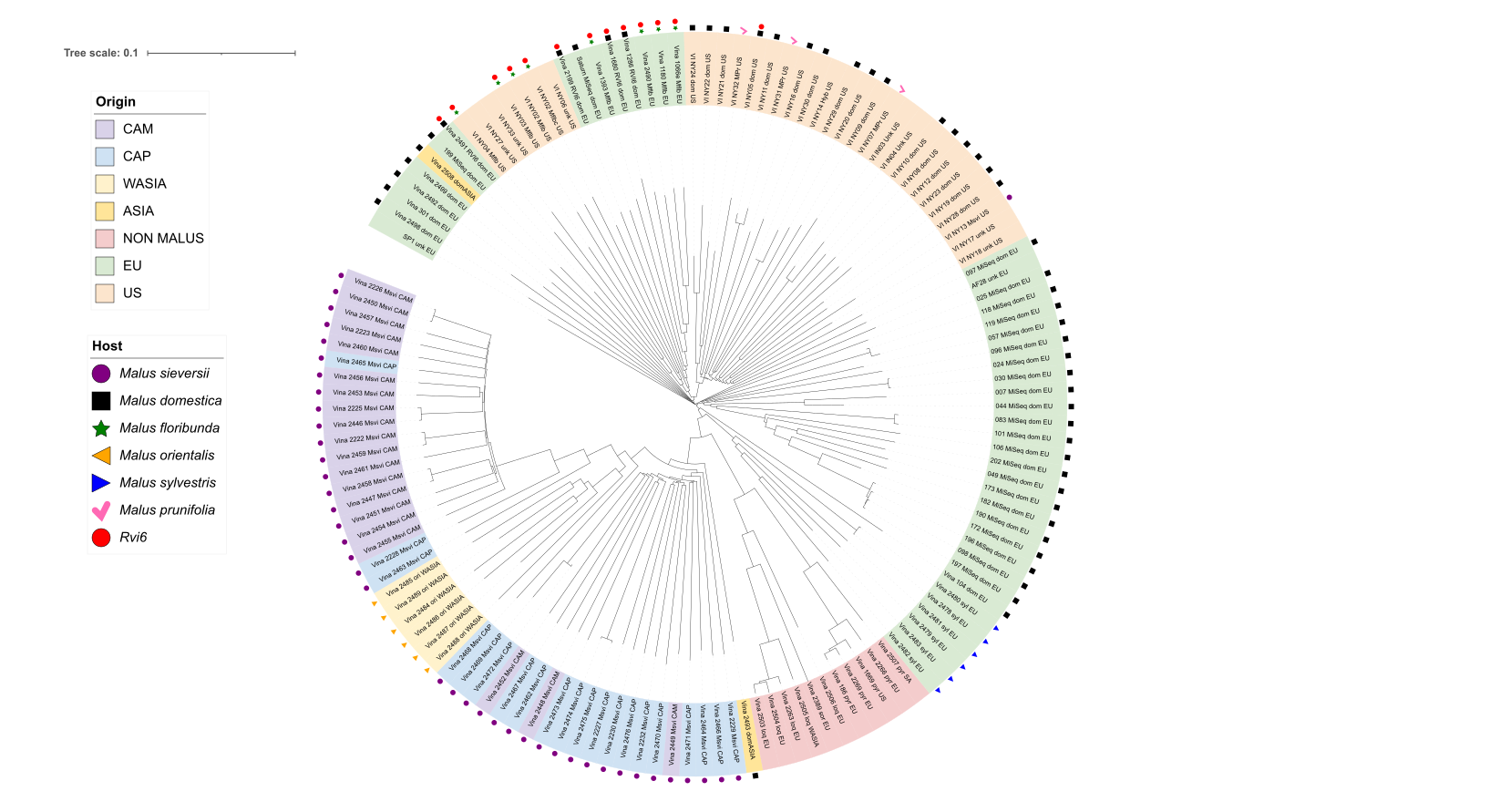

**Supplementary Figure 8.** Neighbor joining tree of *V. inaequalis* from graph-based pangenome derived SNPs, rooted at midpoint. Colored labels indicate geographic origin or if isolates are from a non- *Malus* host. Symbols represent *Malus* hosts if known and isolates collected from hosts with *Rvi6*. Tree visualized using iTOL (https://itol.embl.de/).

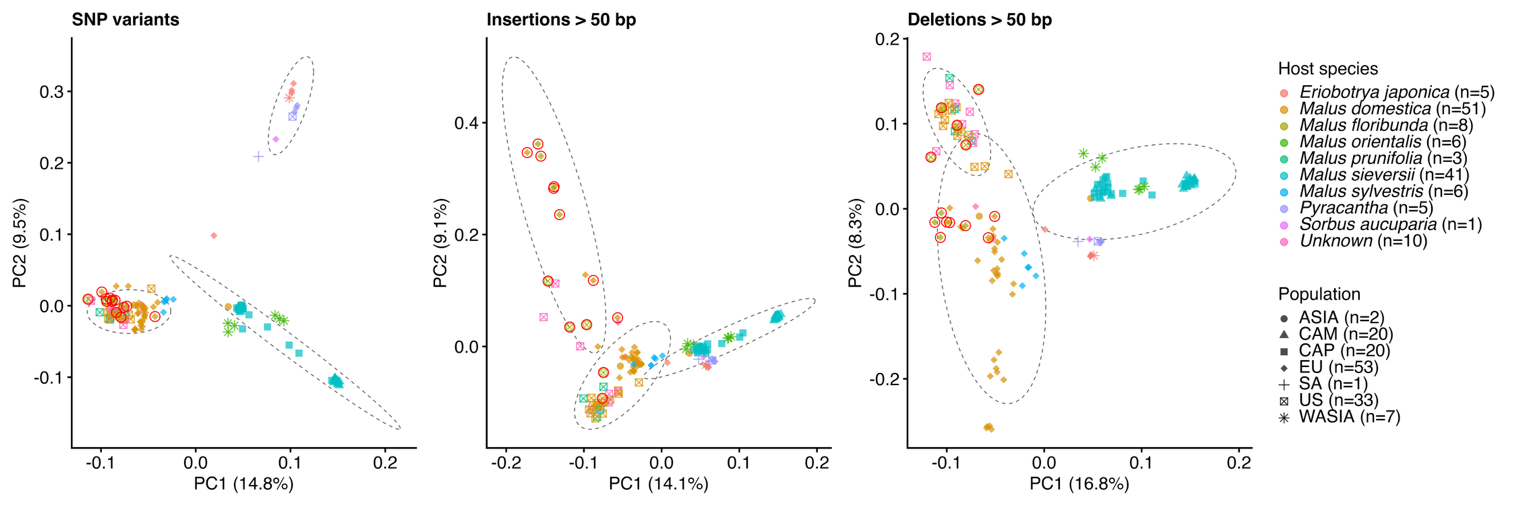

**Supplementary Figure 9.** Principal component analysis of 136 *V. inaequalis* isolates using SNPs**,** Insertions, and Deletions. Dashed ellipses represent k-means clusters identified from PC1 and PC2. Points are colored by the host they were collected from, shapes indicate geographic origin collected from, Red outline indicates the isolate was collected from an *Rvi6* host.

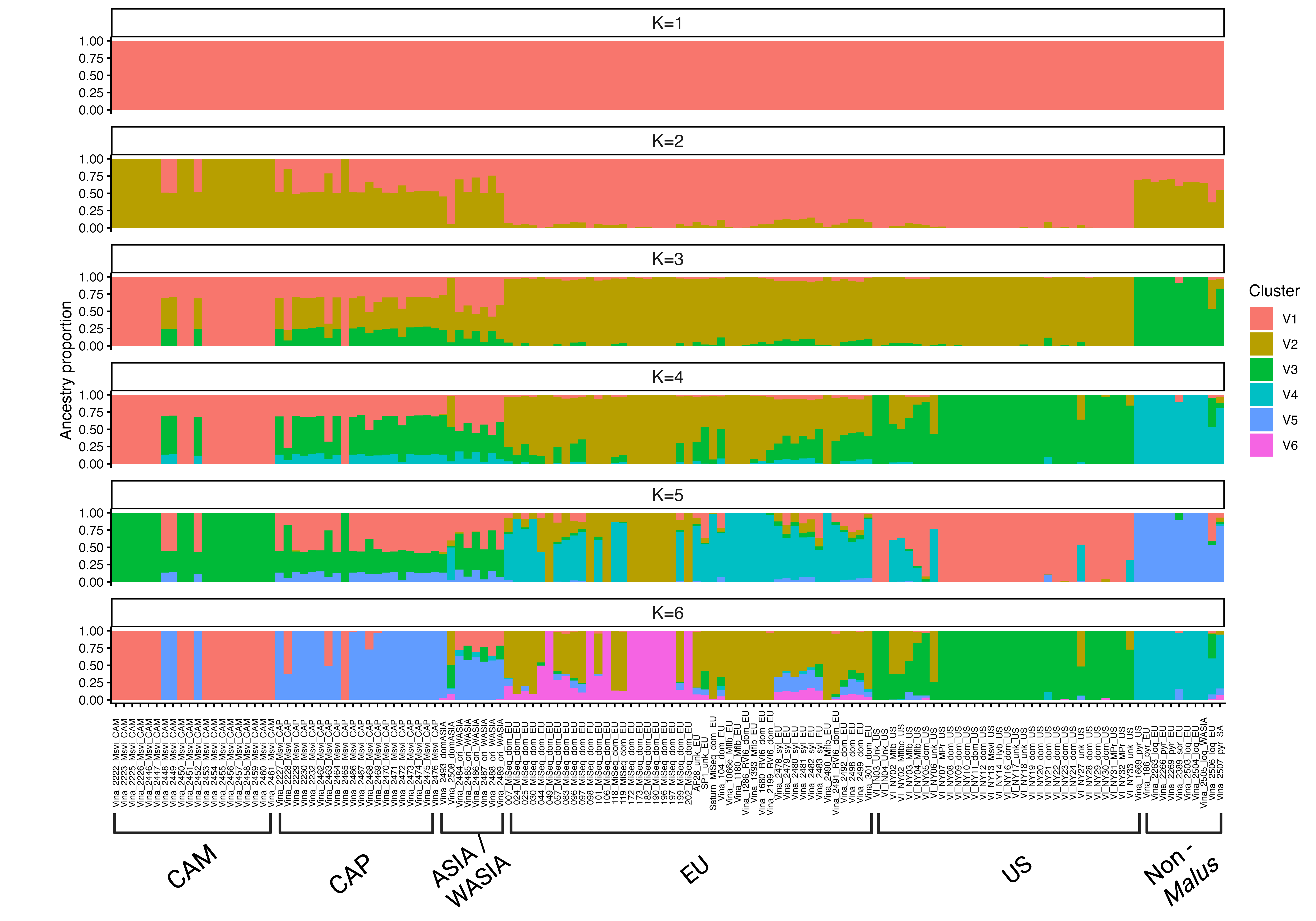

**Supplementary Figure 10**. Population structure of 136 *V. inaequalis* isolates inferred using SNP data, isolate metadata and source in Supplementary Table 16. Isolate populations are listed along the x – axis, CAM (Central Asian Mountains), CAP (Central Asian Plains), Asia and West Asia (China, Japan, and Armenia), Europe, United States, and isolates from non- *Malus* hosts. Colors refer to the ancestral clusters (K 1-6) inferred Admixture v.1.3.0 with SNP genotype data.

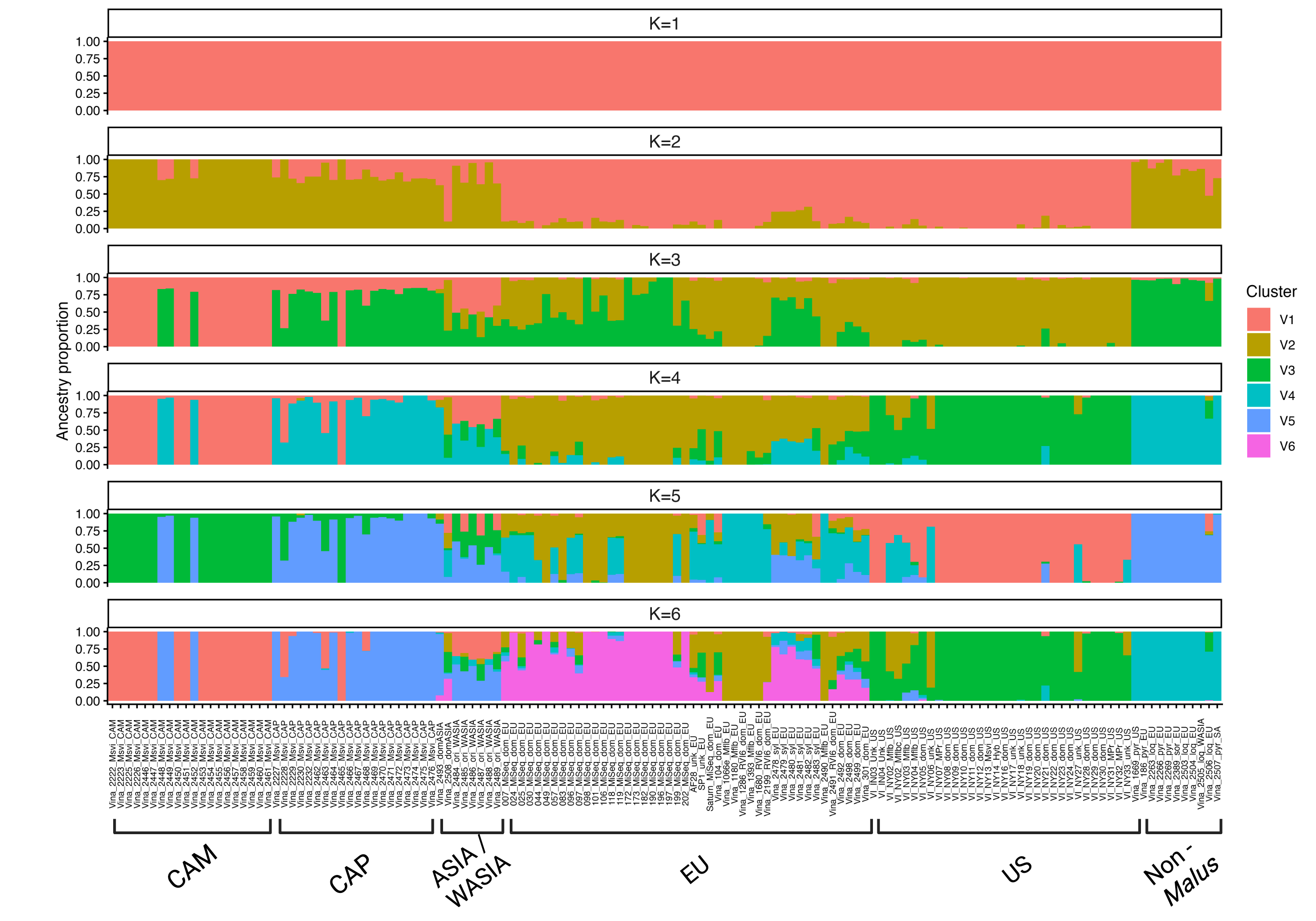

**Supplementary Figure 11**. Population structure of 136 *V. inaequalis* isolates inferred using structural variant data. Isolate populations are listed along the x – axis, CAM (Central Asian Mountains), CAP (Central Asian Plains), Asia and West Asia (China, Japan, and Armenia), Europe, United States, and isolates from non- *Malus* hosts. Colors refer to the ancestral clusters (K 1-6) inferred Admixture v.1.3.0 with structural variant genotype data.

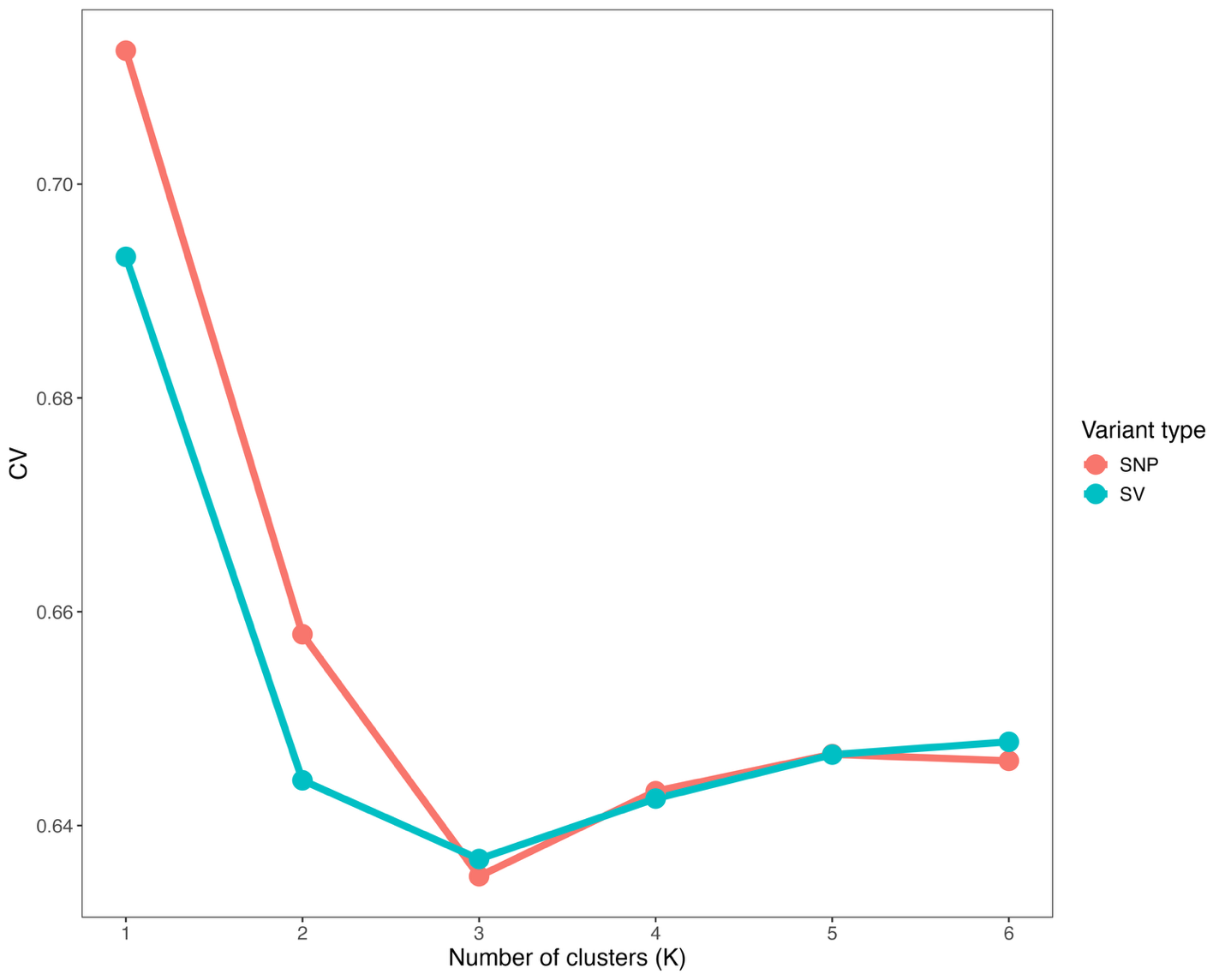

**Supplementary Figure 12.** Cross-validation (CV) error plotted against the number of ancestral clusters (K) from ADMIXTURE v1.3.0 analysis of 136 V. inaequalis isolates. The K value minimizing CV error was selected as the best-supported number of clusters.

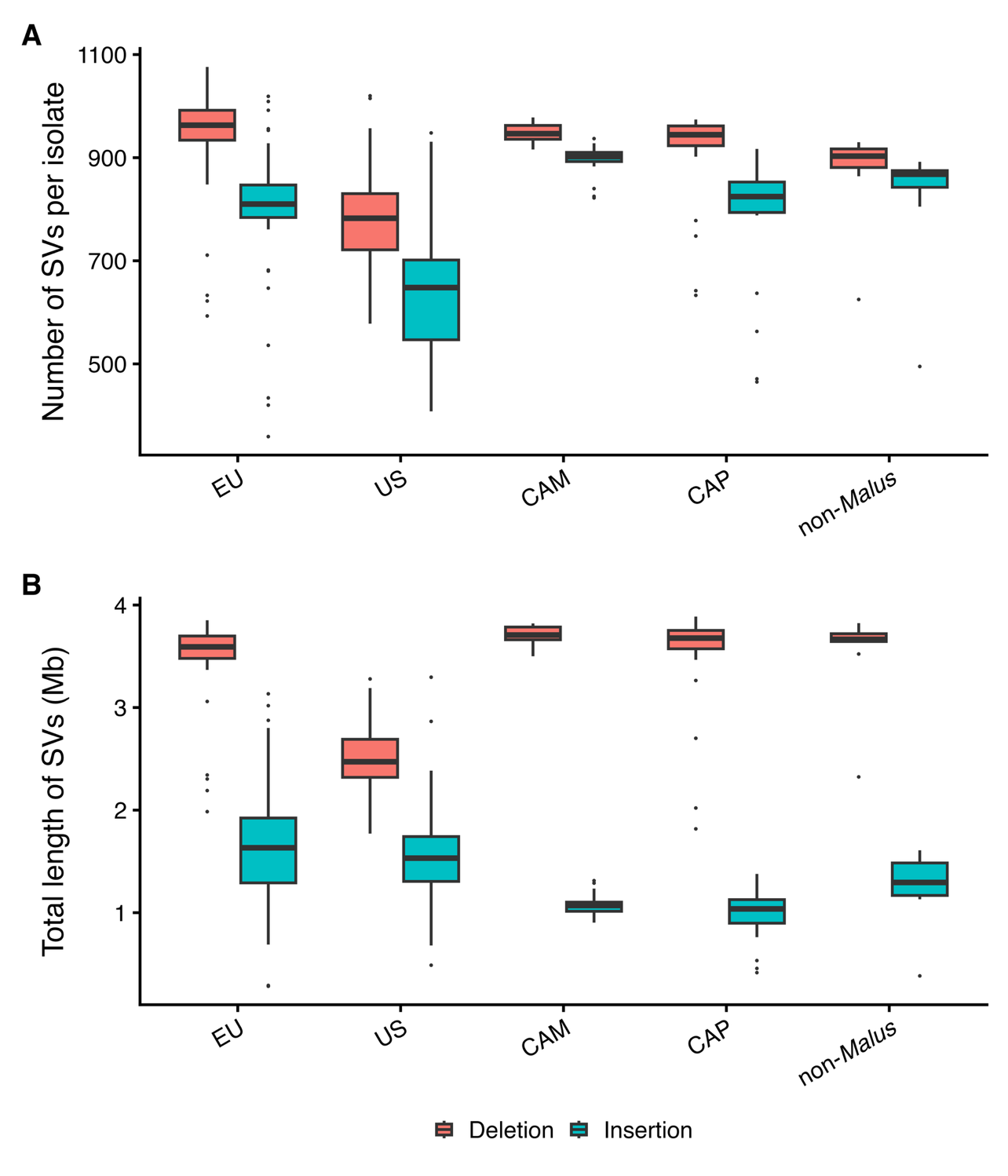

**Supplementary Figure 13.** Structural variant count and length per isolate Boxplots show the distribution of (A) the number of SVs per isolate and (B) the total length of SVs per isolate (Mb) by population of origin: EU (Europe), US (United States), CAM (Central Asian Mountains), CAP (Central Asian Plains), and non-Malus (isolates from non-Malus hosts). Boxes show the interquartile range with the median line; whiskers extend to 1.5 × IQR.

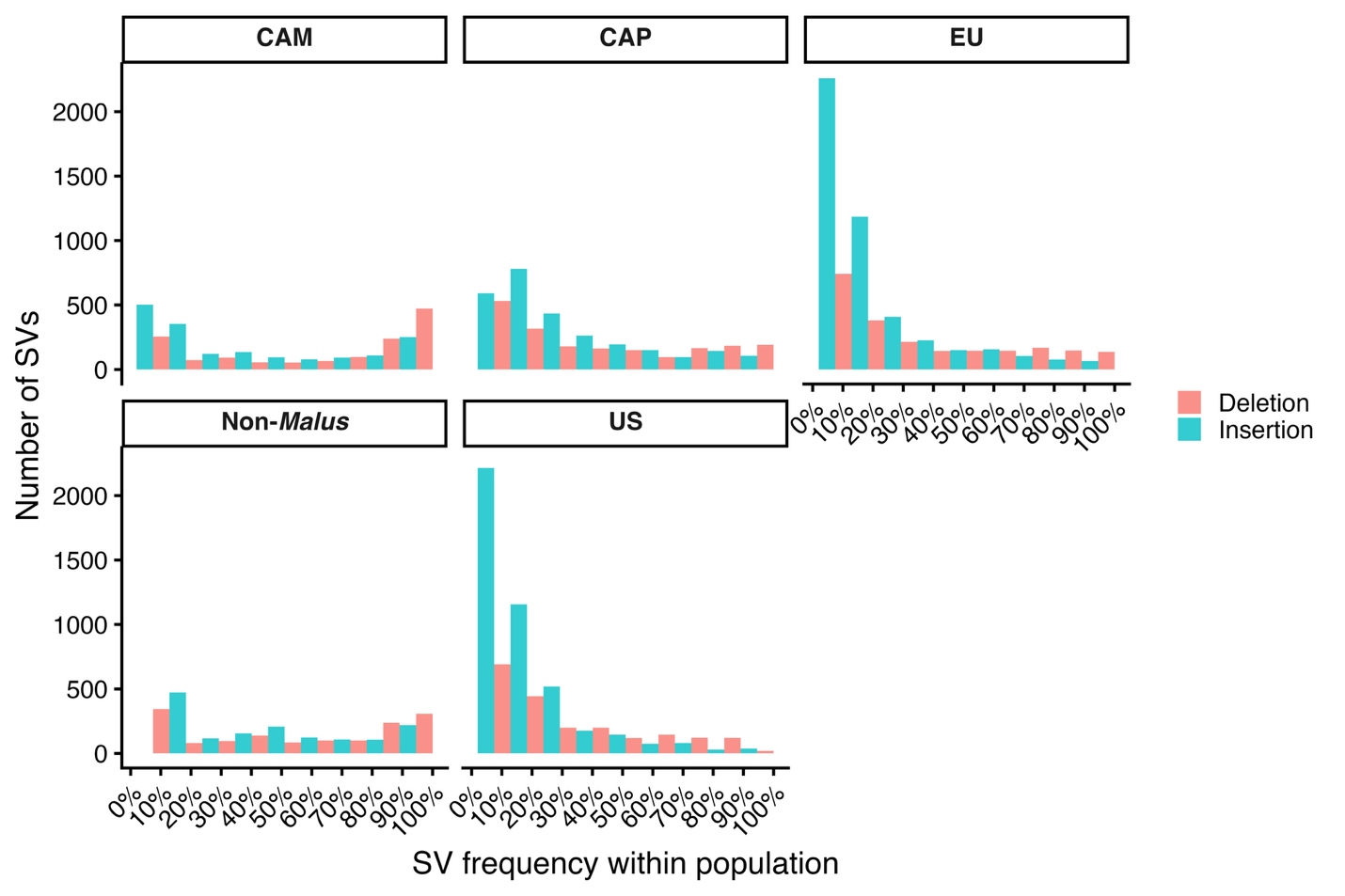

**Supplementary Figure 14. Frequency of structural variants (SVs) within each Venturia inaequalis population, separated by insertions and deletions.** For each population, SVs were binned by their within-population allele frequency (x-axis, 0-100% in 10% intervals), and the number of SVs falling in each frequency bin was counted (y-axis). Panels correspond to populations of origin: CAM (Central Asian Mountains), CAP (Central Asian Plains), EU (Europe), US (United States), and non-Malus (isolates from non-Malus hosts).

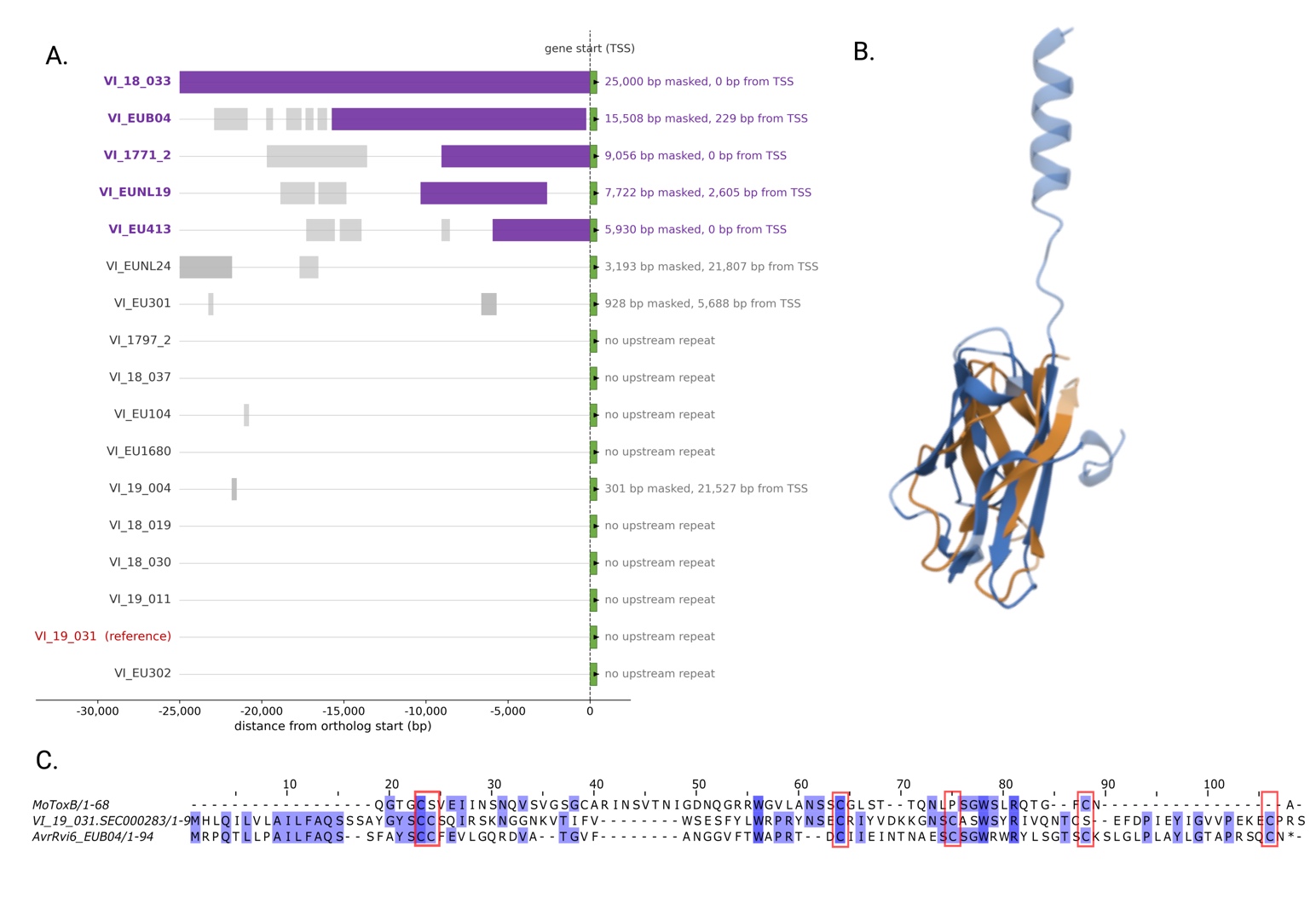

**Supplementary Figure 15**. Analysis of the VI_19_031.SEC000283 MAX- like effector locus in genomes in the *V. inaequalis* pangenome graph. **A)** Repeat content upstream of the orthogroup with VI_19_031.SEC000283. Each row is one isolate, drawn in its own coordinates and aligned at the ortholog start; the green box is the gene start, grey blocks are RepeatMasked (repeat) segments 5′ of the gene. Four assemblies (‘VI-EUB04’, ‘VI-1771-2’, ‘VI-EUNL19’, ‘VI-EU413’) carry a 5.9-15.5 kb repeat block within 0-2.6 kb of the effector start; ‘VI-18-033’ has upstream fully masked, ‘VI-EUNL24’s’ repeat is 21.8 kb upstream; and the reference ‘VI-19-031’ and remaining isolates have non-repetitive upstream sequence. **B)** 3D structural alignment using PDBalign of VI_19_031.SEC000283 (blue) with MoToxB (brown, PDB ID: 6RJ5), Template modeling score (TM-score = .55). **C)** Conserved residues with MoToxB and ArrRvi6^EU-B04^ are shown from ClustalOmega alignment highlighted by similarity, conserved cysteine residues are highlighted.
